# Single-stranded DNA drives σ subunit loading onto RNA polymerase to unlock initiation-competent conformations

**DOI:** 10.1101/2024.08.21.608941

**Authors:** Rishi Kishore Vishwakarma, Nils Marechal, Zakia Morichaud, Mickaël Blaise, Emmanuel Margeat, Konstantin Brodolin

**Author notes:** Rishi Kishore Vishwakarma, Inspiration4 Advanced Research Center Department of Structural Biology, St. Jude Children’s Research Hospital, Memphis, TN 38105, USA.

## Abstract

Initiation of transcription requires the formation of the “open” promoter complex (RPo). For this, the σ subunit of bacterial RNA polymerase (RNAP) binds to the non-template strand of the -10 element sequence of promoters and nucleates DNA unwinding. This is accompanied by a cascade of conformational changes on RNAP the mechanics of which remains elusive. Here, using single-molecule Förster resonance energy transfer and cryo-electron microscopy, we explored the conformational landscape of RNAP from the human pathogen *Mycobacterium tuberculosis* upon binding to a single-stranded DNA fragment that includes the -10 element sequence (-10 ssDNA). We found that like the transcription activator RbpA, -10 ssDNA induced σ subunit loading onto the DNA/RNA channels of RNAP. This triggered RNAP clamp closure and unswiveling that are required for RPo formation and RNA synthesis initiation. Our results reveal a mechanism of ssDNA-guided RNAP maturation and identify the σ subunit as a regulator of RNAP conformational dynamics

## INTRODUCTION

Transcription is the first step in gene expression that is required to interpret the information encoded in duplex DNA. To initiate transcription, all cellular RNA polymerases (RNAPs) must recognize promoter DNA motifs, locally melt duplex DNA and unwind the transcription start site. Duplex DNA is an energetic barrier for transcription initiation and requires the application of mechanical force by RNAPs. In bacteria, transcription initiation is performed by the RNAP holoenzyme (Eσ) assembled from RNAP core (E, subunits α2ωββ’) and the σ subunit that controls promoter recognition and DNA melting. The σ subunits are classified into four groups according to their role and number of structural domains (numbered 1.1 to 4)(1). Group I includes the housekeeping σ subunit (σ^70^ in *Escherichia coli* (*Eco*) and σ^A^ in *Mycobacterium tuberculosis* (*Mtb*) and *Thermus aquaticus* (*Taq*) and controls transcription of most genes during exponential growth. Group II σ subunits (*Eco* σ^38^, *Mtb* σ^B^) have a shorter region 1.1 compared with group I and are implicated in the stress response and stationary phase growth (2, 3). During transcription initiation, σ region 2 (σR2) and region 4 (σR4) recognize the promoter -10 element sequence (T_-12_A_-11_T_-10_A_-9_A_-8_T_-7_) and -35 element sequence (TTGACA) respectively, to form the closed promoter complex (RPc). RPc spontaneously isomerizes to an open complex (RPo) through several structurally distinct intermediates (RPi) (4–6) . In RPo, RNAP is ready to initiate RNA synthesis and forms a stable elongation complex after the synthesis of 11 – 14 nt RNA. The σ subunit regions σR1.1, σR3 and σR4 occupy the DNA and RNA binding channels of the RNAP core and should be displaced in a stepwise fashion by the DNA template (σR1.1 and σR3) and by nascent RNA (σR3 and σR4) during transcription initiation(7).

Binding of σR2 to ssDNA bearing the -10 element sequence (-10 ssDNA) is a key event that triggers promoter melting. σR2 captures the -11A base flipped out of the duplex DNA (8, 9) This results in local DNA melting and then unwinding of ∼13 bp of DNA duplex (8, 10) Base-flipping is the universal mechanism for nucleation of promoter melting by all classes of σ factors classes (11–13). Biochemical studies showed that the free σ subunit and σR2 alone can recognize the -10 motif and bind to -10 ssDNA with low affinity (8, 14). Binding of σR2 to the β’ subunit clamp coiled-coil, called also clamp helices (β’-CH), triggers a cryptic conformational switch in σR2 that increases its affinity for -10 ssDNA (14–17). Although σR2 can bind to -10 ssDNA, neither free σ nor σR2 alone can melt promoter DNA duplex without the RNAP core (18). It has been suggested that the opening-closing dynamics of the RNAP clamp and β subunit lobe (called RNAP pincers) play an essential role in RPo formation by RNAPs in all kingdoms of life (19–22). Single-molecule Förster resonance energy transfer (smFRET) studies of *Eco* RNAP showed that the clamp mainly adopts an open state in the RNAP core and Eσ^70^ holoenzyme and a closed state in RPo where it remains closed during elongation (23–26). In Eσ, the gap between σR2 bound to clamp and β-lobe (10 – 12 Å) is narrower than the diameter of dsDNA helix (20 Å). Therefore, as σR2 obstructs dsDNA access to the RNAP active site cleft (21, 27), the clamp should be opened to allow dsDNA entry and then closed over DNA to hold it in the cleft. Conversely, entry of ssDNA (diameter < 10 Å) does not necessitate clamp opening. The importance of the clamp opening-closing dynamics for RPo formation is supported by studies on antibiotics binding to the RNAP switch regions (28, 29). Specifically, fidaxomicin inhibits transcription initiation by blocking the clamp in the open state (29–32) Conversely, myxopyronin and corallopyronin block the clamp in the closed state (28, 33). Although, clamp dynamics seem to be important for RPo formation, no clear causal relationship has been established between clamp dynamics and promoter melting. Structural studies of promoter melting intermediates formed by *Eco* Eσ^70^ (10, 31) and *Mtb* Eσ^A^ (34, 35) suggest that nucleation of the duplex -10 element DNA melting takes place outside the RNAP cleft and may require clamp closure (31) while entry of the downstream DNA duplex (dwDNA) into the DNA channel requires clamp opening. Unwinding of the transcription start site occurs in the RNAP cleft after dwDNA entry and requires a transient increase in the distance between the β’ switch-2 (β’-SW2) and β fork-loop 2 (β-FL2) that restricts ssDNA access to the active site (34). This scheme fits into the “bind-melt-load-unwind” model and is supported by the results of biochemical studies on real-time RPo formation kinetics (36–38). In the alternative “bind-melt-unwind-load” model, unwinding occurs outside the active site cleft and then ssDNA loads inside (27, 39, 40). Indeed, smFRET studies showed that the clamp remains in the closed conformation during promoter binding and unwinding, suggesting no transient clamp opening during RPo formation (24). Moreover, these models cannot explain biochemical data showing that *Eco* σ^70^R2-R3 and β’ subunit clamp domain (aa 1-314) are sufficient to melt supercoiled DNA duplex (18). As in this “minimal” system clamp closure is irrelevant, it is not clear how DNA unwinding occurs.

In *Mtb*, RPo formation by RNAPs that contain the principal σ^A^ or principal-like σ^B^ subunits is regulated by RNAP-binding protein A (RbpA), essential for bacterial growth (41–44). Our studies depicted RbpA as a transcription factor that stabilizes σ^A^ and σ^B^ interactions with the RNAP core (42, 43), stabilizes RPo, and decreases the energetic barrier for promoter DNA melting (42, 45). RbpA interacts with the β’ subunit Zn^2+^ binding domain (β’-ZBD) and with the non-conserved region (NCR) of σ (σ-NCR) which is regulated by lineage-specific transcription factors (46–48). Biochemical, biophysical and structural studies suggest that RbpA acts as a σ-loader, inducing a σ conformational change that results in σR2 and σR4 stretching over the RNAP core surface to match the distance between the -10 and -35 elements (48, 49). In the absence of RbpA, the Eσ^B^ holoenzyme (but not Eσ^A^) oligomerizes to an octamer in which the RNAP clamp is captured in a fully opened conformation, similar to the one found upon RNAP inhibition with the antibiotic fidaxomicin (50). RbpA binding, which induces σ^B^ remodeling, also leads to clamp closure (30) and octamer dissociation (50)(**Figure 1A**). Here we used *Mtb* Eσ^B^ as a model system to explore the link between clamp dynamics, -10 element recognition, and σ remodeling. We found that the -10 ssDNA fragment is sufficient to induce huge RNAP structure rearrangements, leading to the Eσ^B^ holoenzyme release from a conformational lock. Our results suggest that ssDNA, together with the σ subunit, acts as driver of RPo formation by triggering clamp closure.

**Figure 1.**
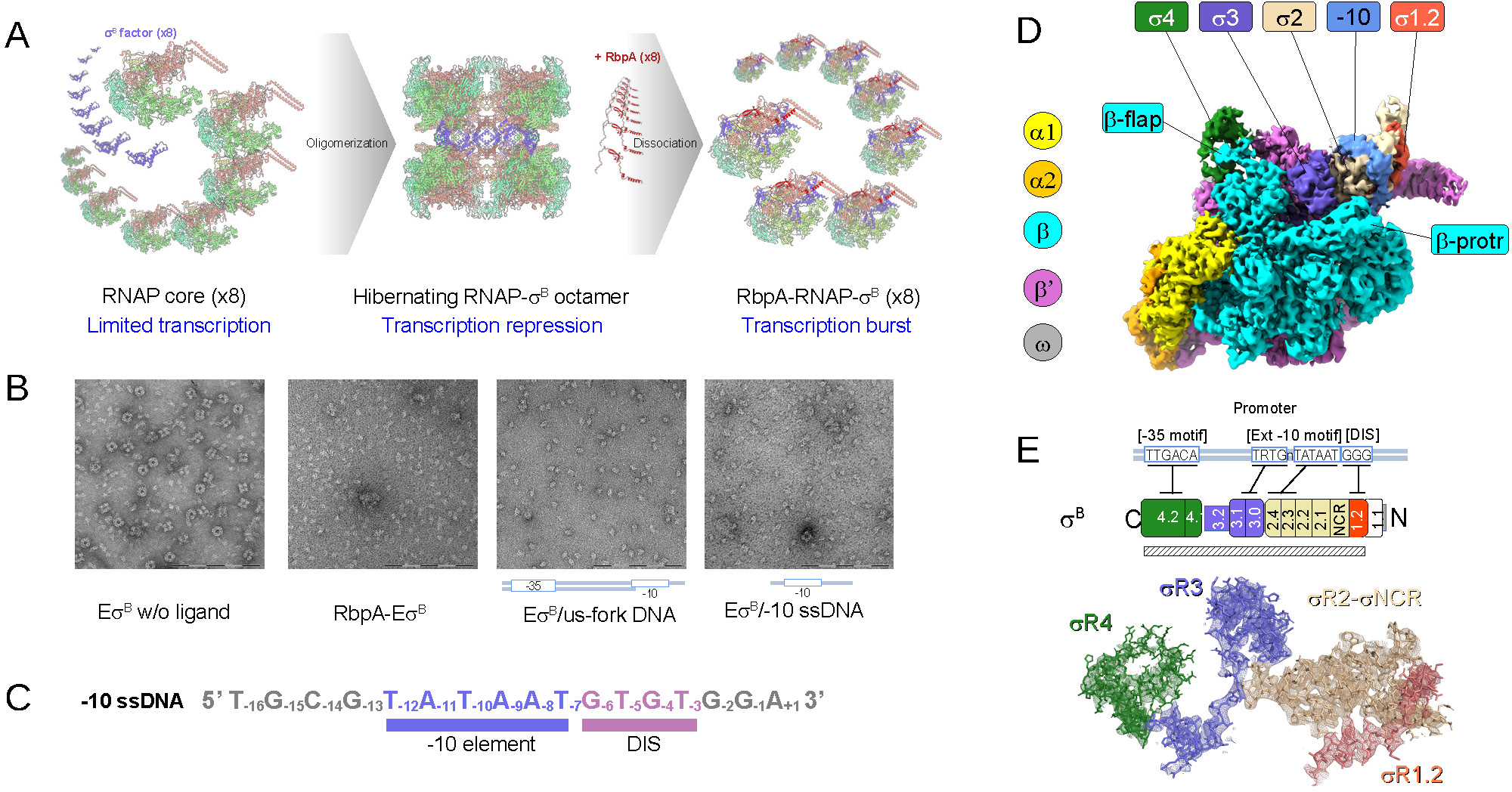
Structure of the *M. tuberculosis* Eσ^B^ holoenzyme in complex with -10 ssDNA. (A) Oligomerization of Eσ^B^ and its regulation by RbpA. (B) Eσ^B^ octamer disassembly induced by RbpA and by -10 ssDNA. Negatively stained images of RNAP holoenzyme octamers without ligand (Eσ^B^ w/o ligand), Eσ^B^ monomers in complex with RbpA (RbpA-Eσ^B^), Eσ^B^ monomers in complex with upstream fork DNA spanning the promoter positions -42/-3 (Eσ^B^/us- fork DNA), and Eσ^B^ in complex with the -10 ssDNA (Eσ^B^/-10 ssDNA). Scale bar = 200 nm. (C) Sequence of the 17-mer DNA oligonucleotide derived from the *lac*UV5 promoter (-10 ssDNA). The locations of the -10 sequence element and discriminator element (DIS) are indicated. (D) Overall cryo-EM map of the Eσ^B^/-10 ssDNA complex (consensus I map). RNAP subunits are color-coded as indicated on the left: yellow and orange for the α subunits, cyan for the β subunit, orchid for the β’ subunit, and gray for the ω subunit. Regions/domains of the σ^B^ subunit are color-coded as indicated on the top: tomato for the σ1.2 region, wheat for the σ2 domain, slate blue for the σ3 domain, forest green for the σ4 domain; and cornflower blue for -10 ssDNA. (E) Cryo-EM density (shown as mesh) and molecular model of the resolved segment of the σ^B^ subunit. The cartoon shows the σ^B^ organization (numbered regions and sub-regions) and its interaction with promoter elements. N-terminus (N) on the right, C-terminus (C) on the left. The subregions inside σ domains are numbered. The hatched rectangle shows the resolved σ^B^ segment (residues 24 -323). The σ^B^ subunit regions/domains σR1.2 (aa 24-55), σNCR (aa 56-86), σR2 (aa 87-163), σR3 (aa 164-240), σR4 (aa 241-323) are color-coded as indicated.

## MATERIAL AND METHODS

### Proteins and DNA templates

The *M. tuberculosis* RNAP core (harboring the C-terminal 6xHis-tag on rpoC) and its mutant (ΔFT), in which amino acids 811-825 of the β subunit were deleted, were expressed in BL21 DE3 *E. coli* cells transformed with the pMR4 plasmid and purified as described before (42, 50). The 6xHis-tagged σ^A^, σ^B^, σ^B^-Cys151/292 mutant, and RbpA were expressed in BL21 DE3 *E. coli* cells and purified as described before (42, 49). Mutant RbpA-R88,89A was constructed using the Quick Change Lightening site-directed mutagenesis kit (Agilent). HPLC-purified DNA oligonucleotides (**Supplementary Table S1**) were purchased from Sigma-Aldrich.

### Protein labeling and smFRET measurements

Random labeling of the σ^B^ subunit with the DY547P1 and DY647P1 fluorescent dyes at Cys151/Cys292 was performed as described before (49). The fluorescent dye derivatives DY547P1-maleimide (donor) and DY647P1-maleimide (acceptor) were purchased from Dyomics GmbH. The double-labeled σ^B^ subunit at 25 pM was prepared in filtered (0.1 µm) FRET buffer [20 mM Tris-HCl (pH 7.9), 150 mM NaCl, 5 mM MgCl_2_, 5% glycerol, and bovine serum albumin (0.1 mg/ml)]. smFRET measurements were performed using a homebuilt confocal PIE-MFD microscope, as described before (49).

### Run-off transcription assays

100 nM of core RNAP and 300 nM of the σ subunit were incubated in transcription buffer (TB) [20 mM tris-HCl (pH 7.9), 50 mM NaCl, 5 mM MgCl_2_, 0.5 mM DTT, 0.1 mM EDTA, 1 µM ZnCl_2_ and 5% glycerol] at 37°C for 10 min. When indicated, RbpA was added to 300 nM. The reaction mixtures were incubated with the *sigA*P and *sigA*P-TGTG derivative (40 nM) promoters at 37°C for 10 min. Transcription was initiated by adding ATP, GTP, UTP to a final concentration of 25 mM/each, 3 μCi of [α-^32^P] CTP (PerkinElmer Life Sciences), 1 mM CTP and was carried out at 37°C for 5 min.

### DNA-protein cross-linking by formaldehyde

Cross-linking reactions and analysis of cross-linked complexes were performed as described before (51) with the following modifications. Briefly, the indicated combinations of RNAP core at 400 nM, σ^A^, σ^B^ at 600 nM, RbpA at 400 nM where mixed in TB and incubated at 37°C for 10 min. The Cy5-labeled DNA oligonucleotide was added to 50 nM and incubated at 22°C for 20 min. Formaldehyde was added to 0.1% and cross-linking performed for 30 sec. Cross-linked complexes were analyzed on 13% SDS-PAGE and gels were scanned with a Typhoon 9400 Imager (GE Healthcare) and stained with Coomassie blue.

### Microfluidics diffusional sizing (MDS) measurements and quantification

4 μM *Mtb* RNAP core and 4.8 μM σ^B^ or σ^A^ and 4.8 μM RbpA (when added) were mixed in 40 μl of binding buffer (20 mM HEPES-KOH pH 8.0, 150 mM KCl, 0.01% BSA, 5% glycerol) incubated at 37°C for 5 min and dialyzed against binding buffer on 0.025μm MF-Millipore membrane filters (VSWP) for 15 min. A series of dilutions were prepared with RNAP concentrations from 4 μM to 15 nM. Then -10 ssDNA labeled by fluorescein at the 5’ end was added at 20 nM final concentration and incubated at room temperature for 20 min. Samples were processed using Fluidity One-W (Fluidic Analytics Ltd). The collected gyration radius values, R_h_ (nm), were normalized and plotted as a radius relative change (*R_RC_*): *R_RC_* = (*R_h_–R_h_0*)/*R_h_0*. *R_h_0* is the gyration radius for DNA alone and *R_h_* is the gyration radius for RNAP-bound DNA. Values from three technical replicates were averaged and fitted using the Grace software (v. 5.1.25) with the Hills equation: *R_FC_ = k[RNAP]^n^/([RNAP]^n^ + K_d_ ^n^)*. *[RNAP]* is the concentration of RNAP holoenzyme, *K*_d_ is dissociation constant, and *k* the amplitude coefficient.

### Cryo-electron microscopy (cryo-EM) sample preparation

15 μM *Mtb*RNAP core was mixed with 22.5 μM σ^B^ in 30 μl of 20 mM HEPES-KOH pH 8.0, 150 mM KCl, 5 mM MgCl_2_, 2mM DTT and incubated at 37°C for 5 min. To remove glycerol traces, samples were dialyzed in 10 μl drops on 0.025 μm MF-Millipore membrane filters (VSWP) against the same buffer at room temperature for 1 h. Then, 3.75 μl of DNA oligonucleotide was added to 22 μM and incubated at 22°C for 30 min. CHAPSO was added to 8mM immediately before sample freezing. About 3.5 μl of sample was spotted on Quantifoil Ultra AuFoil R2/2 Au 200 mesh grids that were prepared using Fischione plasma cleaner (NanoClean model 1070). Grids were flash-frozen in liquid ethane using Vitrobot Mark IV (FEI) at 18 °C and 90% of humidity.

### Cryo-EM data acquisition and processing

Data were collected using a spherical aberration (Cs) - corrected Titan Krios S-FEG instrument (FEI) operating at 300 kV acceleration voltage and equipped with a Gatan K3 Summit direct electron detector (Gatan, Warrendale, PA) and a Gatan BioQuantum energy filter. A total of 9,202 movies (50 frames) were collected at an exposure rate of 27.91 e^-^/Å^-2^/s and total electron dose of 55.735 e^-^/Å^-2^ over a nominal defocus range from -0.8 to -2.5 µm, at a nominal magnification of x81,000 with a physical pixel size of 0.862 Å. Semi-automatic image acquisition was performed with Serial-EM (52). Motion correction, dose weighting, CTF parameters estimation and particles picking were carried out using WARP (53). A set of WARP-selected 695,496 particles was used for further processing. Particles with box size of 360^2^ pixels underwent several 2D classification rounds in cryoSPARC (54). A cleaned dataset of 371,233 particles was used to compute three *ab-initio* 3D models (three classes). The resulting best *ab-initio* model and two junk 3D models were used as references for the 3D heterogeneous refinement and classification. A cleaned set of 290,345 particles from the best 3D class was used for the final local non-uniform refinement resulting in the consensus I cryo-EM map refined to 3.19 Å (**Supplementary Table S2**).

### 3D variability analysis (3DVA) in cryoSPARC

The set of 695,496 particles was re-extracted with box size of 540^2^ pixels and underwent several 2D classification rounds in cryoSPARC. A subset of 394,448 particles from the best 2D classes was used for the *ab-initio* reconstruction to produce five reference volumes: three junk classes, one class corresponding to RNAP dimers, and one class corresponding to RNAP monomers. These volumes were used for the heterogeneous refinement/classification with a larger dataset of 476,952 particles. Next, a subset of 368,449 particles that represented RNAP dimers and monomers was used for the heterogeneous refinement/classification with four reference volumes. The heterogeneous refinement produced two classes of RNAP dimers and one class of RNAP monomers. The final clean dataset of 167,825 particles that represented RNAP monomers was re-extracted with box size of 360^2^ pixels and used in non-uniform refinement to produce the consensus II cryo-EM map refined to 3.33 Å.

#### Separating clamp conformations

A clean dataset of 167,825 particles that included RNAP monomers was used in 3DVA processing with three principal components (reaction coordinates) and a mask excluding σR4. Particles were sorted in three clusters over coordinate 1. The resulting three maps were used as references to classify the set of 167,825 particles using heterogeneous refinement. Class 0, which represents RNAP with the unswiveled clamp conformation (36,319 particles), was refined to 3.79 Å. Class 2, which represents RNAP with the swiveled clamp conformation (21,873 particles), was refined to 4.33 Å.

#### Separating docked and undocked conformations of σR4

The 3DVA processing was repeated with a mask to select σR4. Particles were sorted in four clusters over coordinate 1. The resulting four volumes were used as references to classify the set of 167,825 particles using heterogeneous refinement . Class 1 comprised docked σR4 (72,799 particles) and was refined to 3.43 Å. Class 2 included undocked σR4 (67,957 particles) and was refined to 3.48 Å.

### Model building and refinement

The coordinates of the *Mtb* Eσ^Β^ holoenzyme (PDB ID: 7PP4) were used as starting model. The staring model of full length σ^Β^ was built by AlphaFold. The model of the *lac*UV5 DNA oligonucleotide was built by Coot (55). The molecular models were assembled and fitted to the cryo-EM map using UCSF Chimera (56) and manually modified with Coot. Fitting of the RNAP clamp and lobe domains was adjusted using rigid body real space refinement in Phenix (57). Full cycles of real space refinement in Phenix were performed with secondary-structure restrains and geometry optimization. The refined models were manually adjusted in Coot (**Supplementary Table S2**).

### RNAP conformational heterogeneity analysis in cryoSPARC

3DVA analysis was performed in cryoSPARC with the 167,825 particles using three variability components (eigenvectors). For each component pair, particles were separated in nine clusters and the corresponding cryo-EM maps were calculated. The generated 27 maps and the consensus-I molecular model were used for rigid body real-space refinement with Phenix. (57). Models were aligned with Chimera (56) relative to the RNAP α subunits. Distances between atoms where calculated using custom Python scripts. Clamp rotation was measured with PyMOL Molecular Graphics System using the *draw_rotation_axis.py* script.

## RESULTS

### The promoter -10 ssDNA triggers large scale conformational changes in the Eσ^B^ holoenzyme

Our previous smFRET studies showed that binding of promoter dsDNA to Eσ^B^ does not affect σ^B^ conformation, whereas synthetic upstream fork (us-fork) DNA induces formation of the “open” σ^B^ conformation, like RbpA (49). Here, using negative stain electron microscopy (EM), we observed that the Eσ^B^ octamer disassembled in the presence of us-fork DNA or RbpA (**Figure 1A,B**). We hypothesized that the ssDNA segment of the us-fork comprising the promoter -10 element sequence was responsible for the observed RNAP conformational change. Indeed, addition of a synthetic DNA oligonucleotide (-10 ssDNA) that contains the “perfect consensus” -10 element sequence (T_-12_A_-11_T_-10_A_-9_A_-8_T_-7_) to the Eσ^B^ octamer was sufficient to induce its dissociation (**Figure 1B,C**). To identify the nature of the RNAP conformational changes induced by -10 ssDNA, we determined the structure of the Eσ^B^/-10 ssDNA complex by single-particle cryo-EM to a nominal resolution of 3.2 Å (Consensus I map, **Figure 1D, Supplementary Figure S1, Supplementary Table S2**). In the published structure of the Eσ^B^ octamer, only 44% of the σ^B^ polypeptide was resolved and the cryo-EM density of the σ^B^ subunit C-terminal domains σ3 and σ4 was missing (50). Here, the Eσ^B^/-10 ssDNA complex displayed the complete cryo-EM density of the σ^B^ subunit (93% of σ^B^ resolved) with the C-terminal domain σ3 contacting the β-protrusion, and the domain σ4 inserted into the RNA exit channel and contacting the β-flap (**Figure 1D,E, Supplementary Figure S2A**). The overall σ^B^ fold was identical to that of the primary σ^A^ subunit in the published structures of RNAP-promoter complexes (34, 46), with an root mean square deviation of 1 Å. The cryo-EM densities of domains σ2 and σ3 were well defined (resolution 2.9 – 3.5 Å), while the densities of σR1.1 and σR4 were poorly resolved (resolution: 4 - 6 Å) revealing their high conformational mobility. The central part of the RNAP core, including its active site, displayed the highest resolution (between 2.2 – 3.0 Å) while the mobile domains, β’-clamp with bound ssDNA and β-lobe, displayed a lower resolution (between 2.7 - 5 Å) (**Supplementary Figure S1**). We previously showed that the *Mtb* RNAP core and *Mtb* Eσ^B^ holoenzyme adopt poorly active conformations, characterized by a wide open clamp and a mobile β-flap (50). Superposition of the Eσ^B^ and Eσ^B^/-10 ssDNA structures revealed that the RNAP core displayed large scale rearrangements during holoenzyme maturation (**Figure 2A,B**, **Table 1)**. The clamp domain (β’ residues 4-423, 1219-1261 and β residues 1041-1147), attached to the main RNAP body through switches 1, 2, 3, 5, rotated 22° orthogonal to the main channel and adopted a closed conformation (**Figure 2B**, red). The β-lobe (residues 180-370, blue) rotated 3.59° towards the clamp, bringing the β subunit gate loop (β-GL) close to the σR1.2 and blocking the access to the active site cleft. The β-flap rotated towards the β‘-dock and was fastened by σR4 in a conformation found in all RNAP-promoter complexes structures. In addition, parts of the clamp/jaw/shelf and dock of the β’ subunit (referred as swivel module (58)) exhibited different extents of rotation parallel to the main channel (**Figure 2C,D**). A similar movement, called swiveling, was first observed in the paused elongation complex formed by *Eco* RNAP (*Eco* PEC) (58) and later in *Mtb* PECs (59, 60). The *Mtb* RNAP active site elements (β’ 423-562) that hold the catalytic Mg^2+^ ion and the bridge helix (BH β’ 850-882) also underwent conformational changes upon maturation (**Figure 2D insert)**. Indeed, in the Eσ^B^ octamer, the bridge helix was kinked at β’-A864, adopting the catalysis-inhibited conformation observed in PECs (59). After σR3-σR4 loading, BH adopted a catalysis-ready conformation. Overall, 78% of the β’ subunit and 30% of the β subunit underwent conformational changes. We concluded that the *Mtb* RNAP core and *Mtb* Eσ^B^ holoenzyme adopt a catalytically inactive swiveled conformation, corresponding to an energetically favored, relaxed state. σR3 and σR4 loading onto the RNAP core, induced by -10 ssDNA, forces RNAP to adopt an unswiveled, catalytically active conformation competent to form RPo and initiate RNA synthesis. These results explain the anti-pausing activity of the σR4 in *Eco* σ^70^ and *Mtb* σ^B^ and its capacity to stimulate the initial RNA synthesis (61).

**Figure 2.**
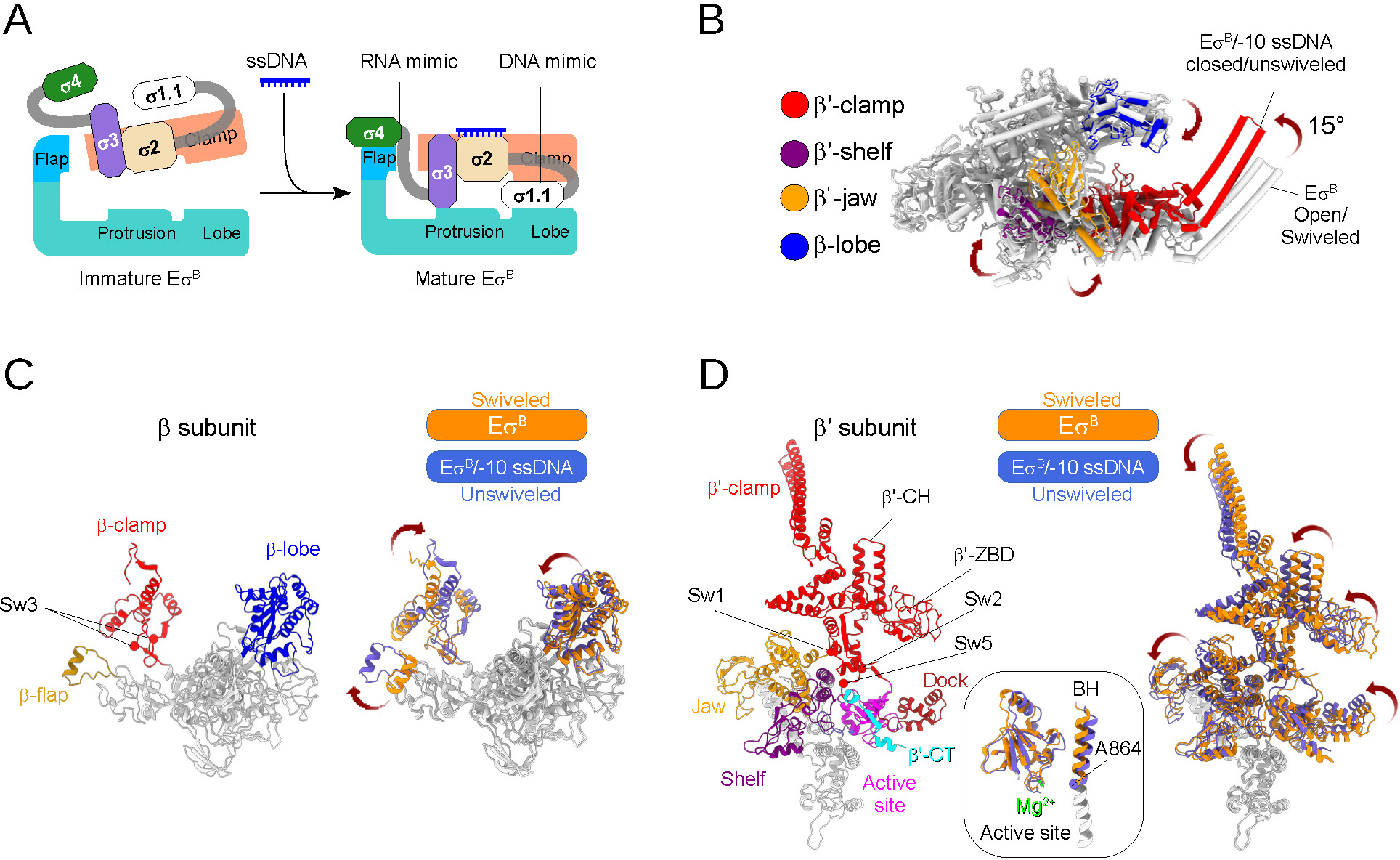
Structural transitions in RNAP during holoenzyme maturation. (A) Cartoon showing the transitions in the Eσ^B^ holoenzyme induced by -10 ssDNA binding. (B) Structure of the Eσ^B^/-10 ssDNA complex superimposed with the structure of the Eσ^B^ octamer (PDB: 7PP4). The RNAP core is shown as light gray ribbons with cylindrical helices. The σ subunit is omitted. The moving domains of Eσ^B^/-10 ssDNA [clamp (β’4 - 419, 1219-1261; β 1117-1140), jaw (β’ 1025-1218), shelf (β’ 882-1011), dock (β’ 444-495), and lobe (β 180-370)] are color-coded as indicated on the left. (C) Ribbon models of the β subunit. Left, static regions are in gray and the moving domains β-clamp (aa 1041-1147), β-lobe (aa 180-370) and β-flap (aa 808-832) are in red, blue and dark goldenrod, respectively. The switch-3 region (Sw3) is delimited by the Cα atoms of β-G1047 and β-G1065. On the right, superposition of the β subunit from the Eσ^B^ octamer (dark orange) with the β subunit from the Eσ^B^/-10 ssDNA complex (slate blue). (D) Ribbon models of the β‘ subunit. Left, static regions are in gray and the moving domains β’-clamp (aa 4-423; 1219-1261) in red, Jaw (aa 1025-1218) in orange, Shelf (aa 882-1011) in purple, Dock (aa 444-495) in firebrick, β‘ C-terminus (β‘-CT, aa 1262-1283) in cyan, active site (aa 423-443; 496-562) in magenta. The σ subunit binding domains, clamp helices β’-CH (aa 339-383) and β’-ZBD (aa 53-85), are indicated. The position of the switch regions is delimited by the Cα atoms of β’-S1219 (Sw1), β’-G419 (Sw2) and β’-P1259 (Sw5). On the right, superposition of the β’ subunit from the Eσ^B^ octamer (dark orange) with the β’ subunit from the Eσ^B^/-10 ssDNA complex (slate blue). Insert shows the changes in the RNAP active site and bridge helix (BH, aa 850-882). The catalytic Mg^2+^ ion is shown in green.

**Table 1.**
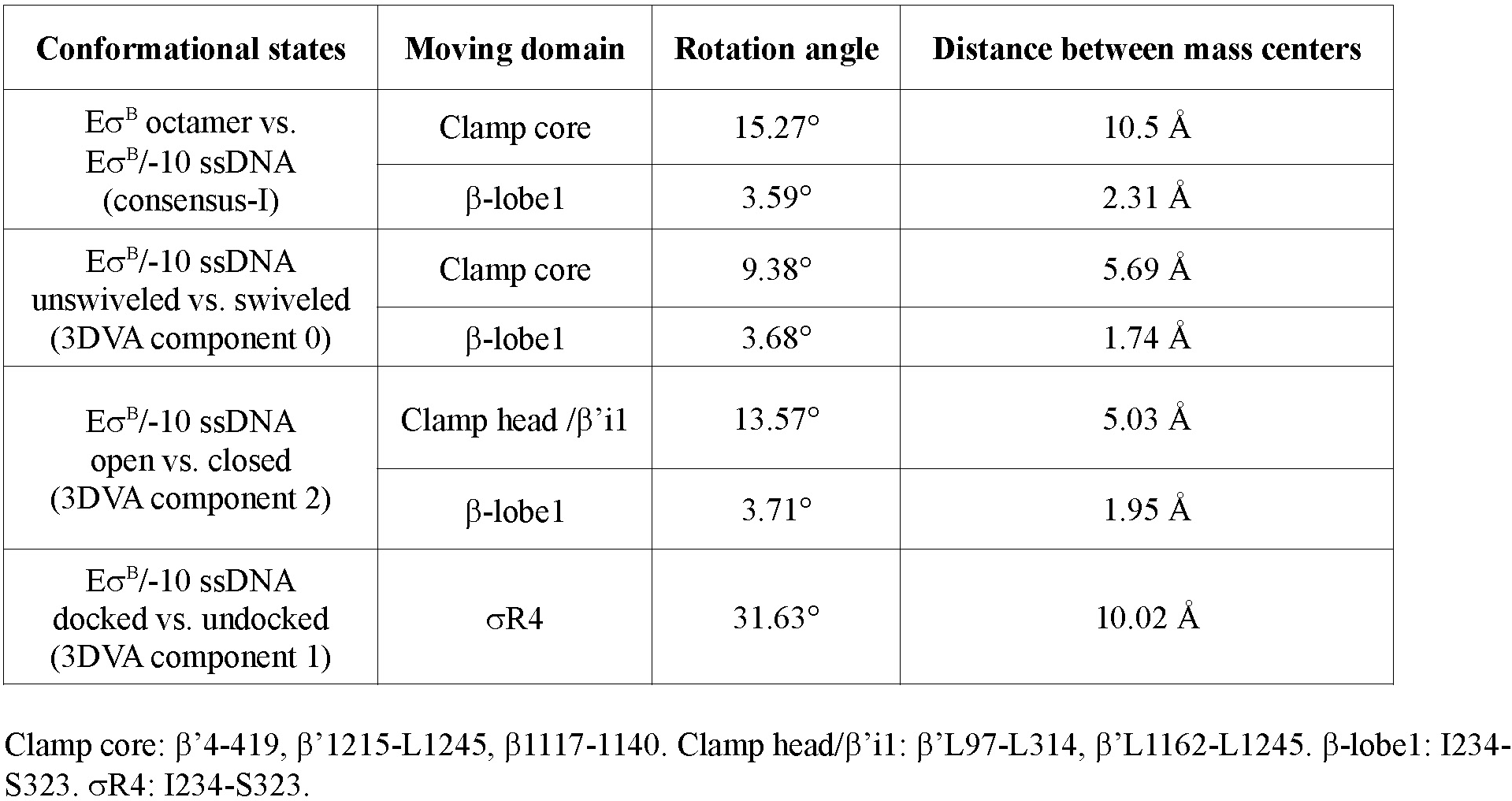
Relative rotation angles of the RNAP domains.

### Changes in the ssDNA-binding interface of σ^B^_2_

The Eσ^B^/-10 ssDNA structure displayed a well-defined cryo-EM density of nucleotides from -13G to -4G with a resolution between 3.0 - 4 Å (**Figure 3A**). This structure included the -10 element (positions -12 to -7) and discriminator element (DIS; positions -6 to -4) (**Figure 3B,C**). The remaining nucleotides were not visible, suggesting that they do not form stable contacts with RNAP. Nucleotides from -13 to -5 interact ed only with the σ^B^ domain that encompasses σR1.2 to sR2 (**Figure 3C**). The -4G of DIS can interact with the conserved residues β-R282 (*Eco* β-R371) and β-E285 (*Eco* β-E374) of β-GL (**Supplementary Figure S2D**). In RPo, when the clamp adopts a fully closed conformation, these residues contact the non-template DNA strand at positions -4, -5 (62). β-GL and β’-CH/σ_2_, direct the non-template DNA strand to the main channel and contribute to RPo stabilization (63). Due to the weak -4G density, we concluded that β-GL does not make stable contacts with ssDNA, but stabilizes binding by sterically restraining fluctuations of the ssDNA 3’-end. The overall path of -10 ssDNA and network of interactions between nucleotides and amino acids (**Figure 3B,C**) matched that of the non-template DNA strand of the transcription bubble in published structures of *Eco* Eσ^70^ RPo (64) and *Mtb* Eσ^A^ promoter melting intermediate RPi2 (34). (**Supplementary Figure S3**). Thus, we concluded that domain σ2 is a major determinant for the path of the non-template DNA strand in RPo.

**Figure 3.**
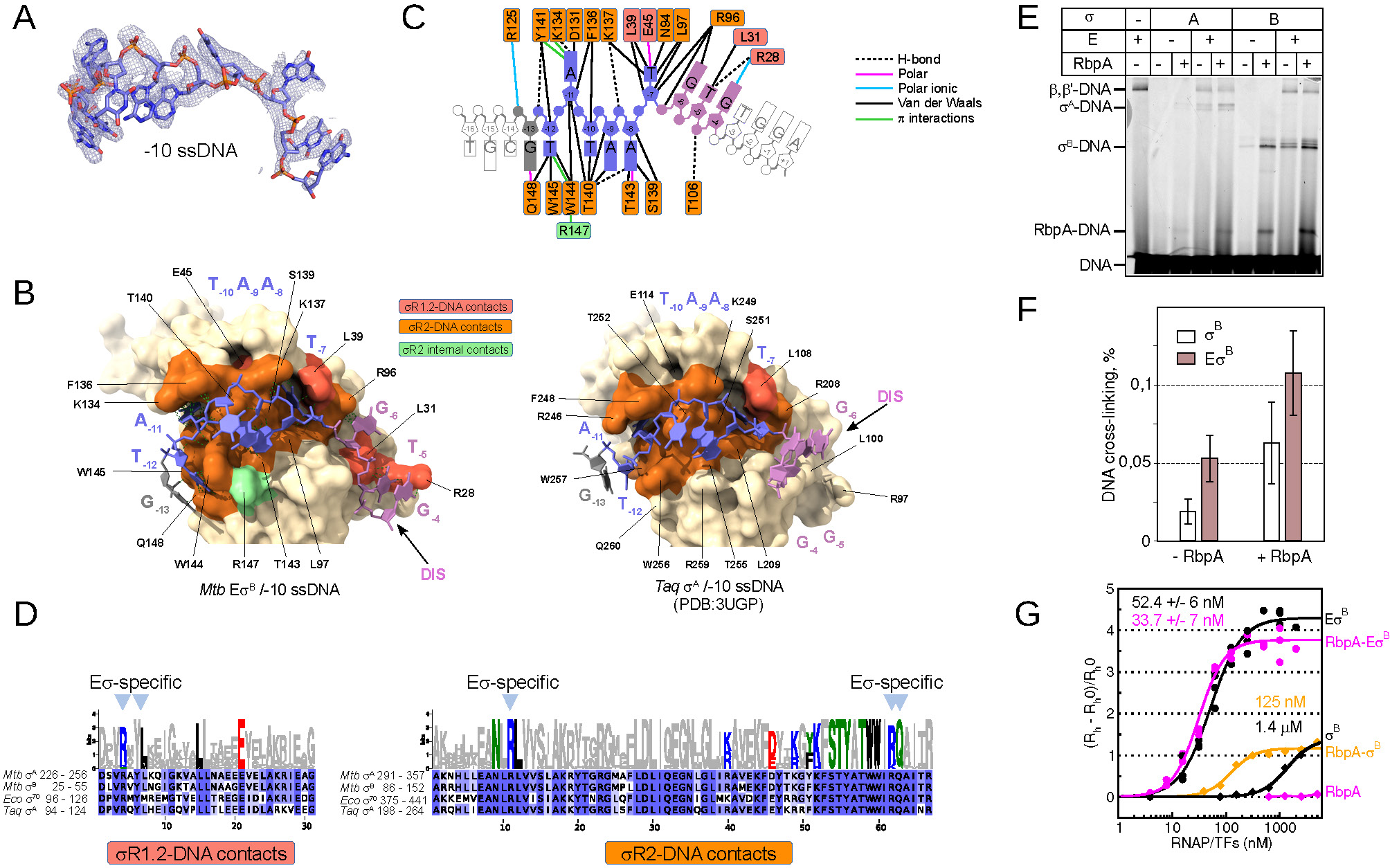
Architecture of the σ^B^/-10 ssDNA interactions and their quantitative analysis. (A) Cryo-EM density (as mesh) and molecular model of -10 ssDNA. (B) Comparison of the σ/ssDNA binding interfaces in the *Mtb* Eσ^B^/-10 ssDNA complex (on the left) and *Taq* σ^A^/-10 ssDNA complex (PDB: 3UGP, on the right). The σ domain 2 is shown as a molecular surface colored in wheat. Residues interacting with ssDNA are in orange (σR2) and tomato (σR1.2). The σ^B^-R147 making holoenzyme-specific π-interactions with σ^B^-W144 is colored in light green. ssDNA is shown as a stick molecular model with filled sugars and bases. Color codes are as in Figure 1E. (C) Schematic drawing showing the network of interactions between DNA and the σ^B^ subunit in the *Mtb* Eσ^B^/-10 ssDNA complex. Resolved DNA bases are color-coded as in Figure 1C. Unresolved bases are in white. The amino acid residues interacting with DNA are indicated and colored as in panel A. Lines are colored according to the interaction type (list on the right). (D) Web logo and alignment of σR1.2 and σR2. The ssDNA contacting residues are colored as follows: polar amino acids (G,S,T,Y,C,Q,N) in green, basic amino acids (K,R,H) in blue, acidic amino acids (D,E) in red, and hydrophobic amino acids (A,V,L,I,P,W,F,M) in black. The arrows on the top indicate the contacts observed in the RNAP holoenzyme and missing in the *Taq* σ^A^/-10 ssDNA complex. (E) Probing of the DNA-protein interactions by formaldehyde cross-linking. *Mtb* RNAP core (E), σ^A^ (A), σ^B^ (B) and/or RbpA (as indicated) were cross-linked to fluorescent -10 ssDNA and resolved on SDS-PAGE. (F) Bar graph showing the quantification of the experiment in panel E. (G) Measurement of the apparent Kd of -10 ssDNA and the σ^B^ subunit by MDS. The relative change *RRC* of the mean Rh is plotted in function of the protein concentration. The calculated values of the apparent Kd are shown.

To understand how the RNAP core stimulates -10 ssDNA binding to σ2, we compared the Eσ^B^/-10 ssDNA structure with the published crystal structure of the *Taq* σ^A^2-3 fragment in complex with -10 ssDNA (8) (**Figure 3B**). The overall path of the -10 element nucleotides (blue) was identical in the two structures. However, the orientations of residues -6 to – - 4 in DIS (pink) and of the upstream “fork region” residues -13 to -12 (gray) were different. In the *Mtb* Eσ^B^/-10 ssDNA complex, two DIS nucleotides (-5T and -6G) contacted the groove formed by σR1.2 (residues σ^B^-R28, σ^B^-L31) and σR2.1 (σ^B^-R96). In the *Taq* σ^A^2-3/-10 ssDNA structure, these interactions were missing because DIS nucleotides were displaced outside the σR1.2 groove. The lack of interaction with DIS explains the difference in affinities to -10 ssDNA displayed by the free σ subunit and RNAP holoenzyme reported earlier (14, 17) and see below. Concerning the us-fork region, -12T interacted with σ^B^-Q148 (*Eco* σ^70^-Q437, *Taq* σ^A^ -Q260). The -12T nucleotide, which is recognized as a base pair in RPo; was unpaired in the *Mtb* Eσ^B^/-10 ssDNA complex analogously to the melting intermediate complex T-RPi1 (10) . In the Eσ^B^/-10 ssDNA complex, -12 T interacted with the invariant σ^B^-W144,145 residues of σR2.3 (W-dyad, *Eco* σ^70^-W433,434, *Taq* σ^A^ -W256,257). The W-dyad was stabilized in a “chair-like” conformation through π-stacking with σ^B^-R147 (**Figure 3C, Supplementary Figure S2D**). The W-dyad chair-like conformation is characteristic of RPo (**Supplementary Figure S2F)**. W-dyad isomerization from the edge-on to the chair-like conformation occurs after nucleation of -11A melting, but before melting of +1, and correlates with the transcription bubble propagation from -11 up to -9 (10). In T-RPi1, -11A was flipped out, but W-dyad was still in the edge-on conformation (**Supplementary Figure S2E**). In the *Taq* σ^A^2-3/-10 ssDNA structure, -12T was displaced and could not interact with *Taq* σ^A^ -Q260 (σ^B^-Q148) and W-dyad. The W-dyad of the *Taq* σ^A^2-3/-10 ssDNA complex was in the edge-on conformation, like in free σ^70^ (65) and in RNAP holoenzymes (**Supplementary Figure S2A,B,C**). Therefore, we suggest that binding of the ssDNA segment between -11 to –5 to σ forces W-dyad isomerization into the chair-like conformation that is further locked by interaction with σ^B^-R147 and stacking with -12T. As we did not observe this RPo-specific conformation in the *Taq* σ^A^2-3 apo complex, we suggest that β-GL stabilizes the interaction of σR1.2 with DIS and thus in turn stabilizes upstream ssDNA contacts. Functional studies demonstrated that *Taq* σ^A^-W256 (σ^B^-W144) is not required for -10 ssDNA binding, but is implicated in RPi isomerization to RPo (66). Therefore, ssDNA binding to domain σ2 acts as a trigger for isomerization in the context of the RNAP holoenzyme. To conclude: (1) σ binding to the RNAP core does not change the ssDNA-binding interface of domain σ2; (2) -10 ssDNA binding to the RNAP holoenzyme leads to changes in the ssDNA-binding interface of domain σ2, such as W-dyad isomerization and contacts formation between σR 1.2 and the DIS element. Contacts with DIS might be stabilized by the interactions with β-GL. These findings explain a large set of biochemical data showing that the RNAP core stimulates -10 ssDNA binding to σ2 (14, 16, 67).

### The RNAP core and RbpA stabilize the interaction of the σ_2_ domain with -10 ssDNA

*Eco* σ^70^ and *Taq* σ^A^ models have been used to investigate how -10 ssDNA binding to region 2 of the σ factor is stimulated by the RNAP core. However σ^70^ and σ^A^ differ from *Mtb* σ^A^/σ^B^ by the presence of large insertions in their NCR. These insertions could partly explain the observed “stimulation”. To determine whether the *Mtb* RNAP core also stimulates *Mtb* σ^A^/σ^B^ binding to -10 ssDNA, we used a formaldehyde cross-linking assay that gives a relative estimation of protein-ssDNA affinity (51, 68). After cross-linking the RNAP core, free σ^B^ subunit and Eσ^B^ to -10 ssDNA (end-labeled with Cy5) in the presence or absence of RbpA, we analyzed the obtained DNA-protein complexes by SDS-PAGE followed by Cy5 fluorescence imaging (**Figure 3E)** or Coomassie blue staining (**Supplementary Figure S4A**). In agreement with previous findings using *Eco* Eσ^70^ (14, 16), the *Mtb* RNAP core produced one slow-migrating cross-linked species with a mobility that corresponded to the β/β’ subunits. Conversely, we observed only a weak cross-linking between DNA and the free σ^B^ subunit. In the presence of RbpA, we detected a fast-migrating band at the bottom of the gel, above the free DNA that we assigned to RbpA-DNA cross-linking (marked as “RbpA-DNA”). The σ-specific cross-linking was increased by ∼2 fold in the presence of the *Mtb* Eσ^B^ holoenzyme compared with free σ^B^ (**Figure 3E,F)**. This indicated that the, *Mtb* RNAP core stimulateed -10 ssDNA binding to the σ^B^ subunit, like the *Eco* RNAP core to σ^70^. Addition of RbpA to free σ^B^ resulted in a similar level of σ^B^-DNA interaction stimulation, as observed with the RNAP holoenzyme alone (∼3-fold higher stimulation relative to free σ), indicating similar affinities of -10 ssDNA for the σ^B^-RbpA complex and the *Mtb* Eσ^B^ holoenzyme. The combination of σ^B^, RbpA and *Mtb* RNAP core resulted in a cooperative effect that was reflected by the further increase of σ-specific cross-linking. We concluded that both RbpA and RNAP core stabilize the interaction of σ^Β^_2_ with the -10 element. Cross-linking experiments with the σ^A^ subunit demonstrated that RbpA also stimulated binding of -10 ssDNA to free σ^A^ and to the Eσ^A^ holoenzyme to a similar extent as observed with σ^Β^ (**Figure 3E**). RbpA stimulated σ^B^-DNA cross-linking even in the presence of a truncated -10 ssDNA (positions -12 to +1) that lacked the RbpA-contacting nucleotides -14/-15 (**Supplementary Figure S4B**). Conversely, Hubin et al., (46) reported that RbpA does not affect binding of truncated -10 ssDNA to the σ^A^ subunit in the context of the *Mtb* RNAP holoenzyme. The origin of this discrepancy remains unclear.

To quantify the interactions between the σ subunit and -10 ssDNA, we used MDS to measure the average gyration radius (*R_h_*) of diffusing macromolecules labeled with a fluorescent probe. To decrease possible artifacts from protein aggregation, we used a high ionic strength buffer (150 mM KCl) without Mg ^2+^ ions. The *R_h_* value of -10 ssDNA end-labeled with fluorescein was ∼1.9 nm. Addition of Eσ^Β^ increased the R_h_ value to ∼10 nm (**Supplementary Figure S4C**). We did not observe any significant change using an oligonucleotide that carried a substitution of the invariant “master” base A_-11_ to C in the -10 motif (-11C ssDNA) to abolish the interaction between σR2 and the -10 element (8, 9, 69) (**Supplementary Figure S4C**). Therefore, we concluded that the MDS assay detected specific interactions between -10 ssDNA and RNAP. Next, we measured the affinity for -10 ssDNA of σ^Β^ alone, RbpA-σ^Β^ complex, Eσ^Β^ and RbpA-Eσ^Β^ (**Figure 3G**). We presented data as the relative change in gyration radius (*R_RC_*), calculated as the change in *R_h_* at a given protein concentration divided by the *R_h_0* of DNA alone: *R_RC_*=(*R_h_ - R_h_0*)/*R_h_0*. RbpA alone did not induce any detectable *R_h_* change, indicating that it does form a stable complex with ssDNA. The σ^Β^ subunit displayed low affinity for -10 ssDNA (apparent K_d_ of 1.4 μM). In the presence of RbpA, σ^Β^ affinity for -10 ssDNA increased by ∼11-fold (K_d_ ∼ 125 nM). In the presence of the Eσ^Β^ holoenzyme, affinity increased by 27-fold (K_d_ ∼ 52 nM). Addition of RbpA to the Eσ^Β^ holoenzyme further increased the affinity by ∼1.6-fold (K_d_ ∼ 34 nM), in agreement with the results of the formaldehyde cross-linking assay. Altogether, these experiments showed that binding to σ^Β^ of RbpA or RNAP core stabilizes its interaction with -10 ssDNA.

### RNAP in complex with -10 ssDNA adopts multiple conformational states

The consensus I cryo-EM map of the *Mtb* Eσ^B^/-10 ssDNA complex revealed the conformational mobility of the RNAP clamp and σ^Β^R4. To find the range of conformational states adopted by RNAP in our sample, we performed a 3DVA using cryoSPARC (54, 70). The 2D-aligned images of the Eσ^B^/-10 ssDNA complex revealed two particle populations: RNAP monomers and RNAP dimers (**Supplementary Figure S5, 2D-classification**). For the 3DVA, we selected a subset of particles that included only RNAP monomers. First, we refined the Eσ^B^/-10 ssDNA monomer map (consensus-II map) to a nominal resolution of 3.3 Å (**Supplementary Figure S5)**. The structure of this consensus II Eσ^B^/-10 ssDNA complex was identical to that of the consensus I map. Next, we performed 3DVA on the consensus II map using three variability components (eigenvectors) **(Supplementary Figure S6)**. Reconstruction of a series of twenty intermediate maps over each 3DVA component revealed a full range of RNAP conformational states (**Supplementary Figure S6C, Movie 1**). The first and the last intermediates represented the boundary states with a maximum amplitude in domain movement relative to the “average” consensus structure (**Table 1**). We observed three conformational change types: i) swiveling of the clamp core (β’ residues 4-419, 1215 -1245, **Figure 4A, pink**) parallel to the main cleft with simultaneous closing, orthogonal to the main cleft (components 0 and 2); ii) closing of the clamp head (β’ residues 97-314, 1162-1245, **Figure 4B, pink**), orthogonal to the main cleft (component 0 and 2); and iii) swinging of the σR4/β-flap module between the β’-dock and β’ Zn^2+^-binding domain (β’-ZBD) (component 1) (**Figure 4D**). In components 0 and 2, clamp movement was accompanied by β-lobe closing (residues 180-370, **Figure 4A,B blue**). Clamp and lobe movements were gradual, while σR4 hopped between boundary states (**Supplementary Figure S7, S8**).

**Figure 4.**
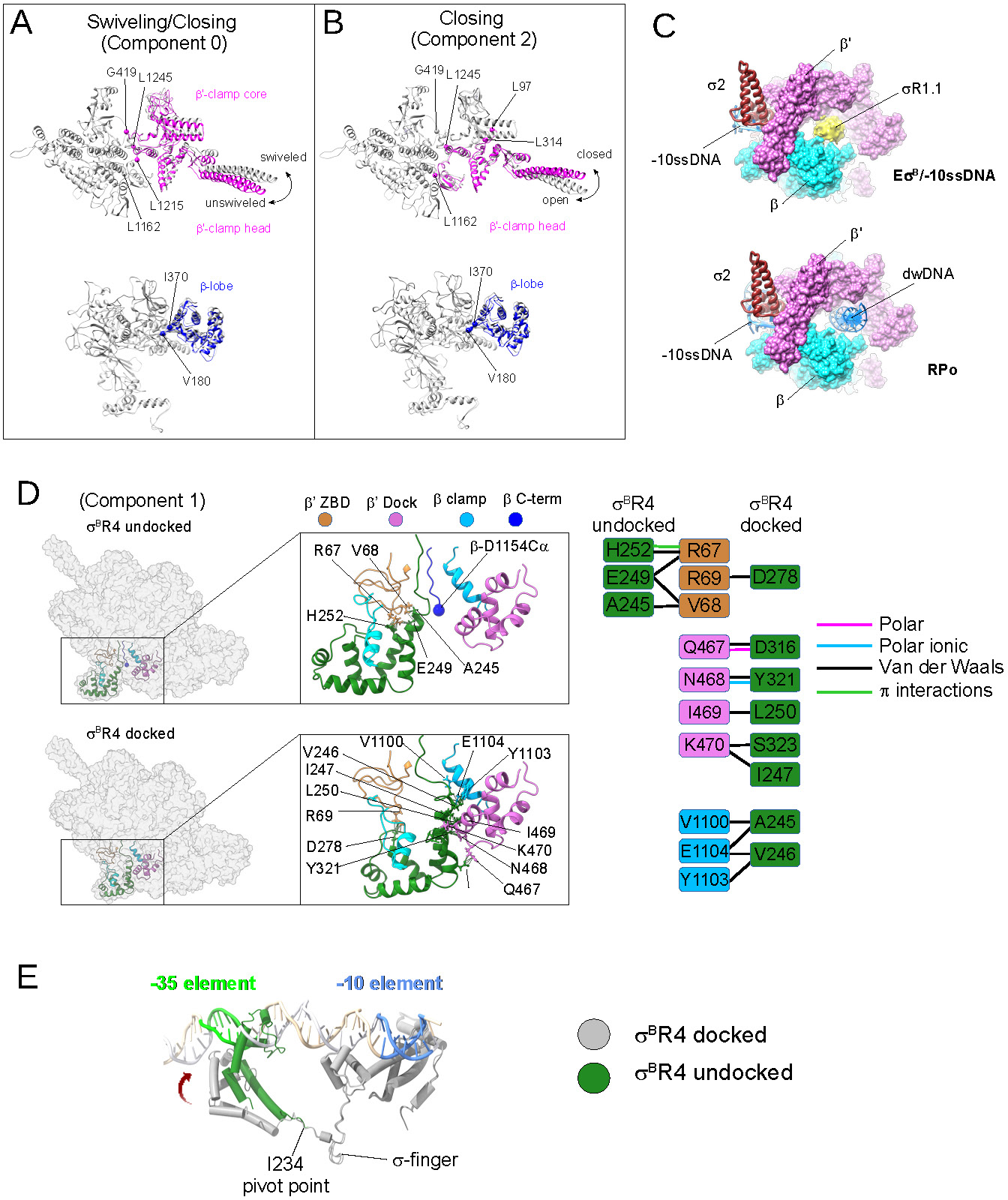
Conformational landscape of Eσ^B^ bound to -10ssDNA. (A) Conformational changes in the RNAP core for the 3DVA component 0: clamp swiveling and closing. The β and β’ subunit domains that underwent conformational changes are in blue and pink, respectively. (B) Conformational changes in the RNAP core for 3DVA component 2: clamp closing. The β and β’ subunit domains that underwent conformational changes are in blue and pink, respectively. (C) Comparison of the position of σR1.1 in the unswiveled Eσ^B^/-10ss DNA complex and position of the dwDNA in RPo. RNAP is shown as a molecular surface color-coded as in Figure 1D. The σ^B^ subunit is shown as ribbons. The cryo-EM density of σR1.1 is represented as a yellow surface. (D) Conformational changes for the 3DVA component 1: σR4 swinging between the docked and undocked state. RNAP is represented as a molecular surface. The σR4 and RNAP core domains (β’-dock, β’-ZBD, β-clamp) contacting σR4 are shown as ribbon models. The contacting residues are shown as ball & stick molecular models. The cartoon on the right shows the map of the interactions between σR4 and RNAP core in the undocked and docked σR4 states. Lines are colored according to the interaction type. (D) Superposition of the σ^B^ subunit in the docked (gray) and undocked (green) conformations with the promoter DNA structure from the *Mtb* RPc (PDB: 7KIM).

### Region 1.1 of **σ^B^** occupies the downstream DNA channel when the clamp is closed and unswiveled

To determine the structures of the most populated boundary states of the clamp, we refined the maps and sorted particles according to the 3DVA (**Supplementary Figure S8**). We computed two Eσ^B^ structures that corresponded to boundary clamp states: swiveled clamp (swRNAP) and unswiveled clamp (usRNAP). For both states, the -10 ssDNA conformation was identical except its 3’ end nucleotides -4/-5 that fluctuated between β-GL and β protrusion. The most noticeable change was in the cryo-EM density assigned to σ^B^R1.1 (residues 1-23) and located in the DNA-binding channel (σR1.1, **Figure 4C**). Previously we showed that in the *Mtb* Eσ^B^ holoenzyme, residues L17-A24 of σ^B^R1.1 were stacked to the clamp head surface, while the cryo-EM density of σ^B^ N-terminus was missing. In usRNAP, the missing density of σ^B^ R1.1 was well defined and occupied the place of the downstream promoter DNA duplex (dwDNA, **Figure 4C**). We concluded that in the *Mtb* Eσ^B^ holoenzyme, σ^B^R1.1 stays in the RNAP DNA-binding channel, but adopts multiple conformational states or unfolds. In usRNAP, σ^B^R1.1 movements are restrained by the clamp/lobe and therefore, its density become better defined. We propose that transient clamp opening weakens the interactions of σ^Β^R1.1 with RNAP and allows dwDNA to displace the σ^Β^R1.1 out of the DNA channel, an essential step for the isomerization from the RPi to RPo.

### Docking of **σ**R4 in the RNA channel coincides with the β subunit C-terminus restructuring

To determine the structures of the most populated boundary states of σR4 we performed 3DVA with a mask on σR4 and calculated maps from the particles sorted according to component 1 (**Supplementary Figure S6, Figure S8**). Particles were distributed equally between two distinct RNAP conformations (43.4% of docked σR4 and 40.5% of undocked σR4). The respective maps were refined to 3.43 Å and 3.48 Å. In the RNAP bearing docked σR4, the overall σ^Β^ path matched that of σ^Β^ in the consensus I/II structures and that of σ^A^ in the published structures of *Mtb* RPo. Moreover, σR4 was inserted in the RNA-exit channel and its residues σ^Β^-I247, L250, D316, Y321 and S323 contacted the β’-dock domain, residue σ^Β^-D278 contacted the β’-ZBD, and residues σ^Β^-A245 and V246 contacted the β-clamp (**Figure 4D**). The structure of RNAP with undocked σR4 was different from all published RNAP holoenzyme structures. First, the RNA exit channel was partially occluded by the C-terminal tail (CTT) of the β subunit (residues 1148-1154) that inserts between the β’-ZBD and the β-clamp (**Figure 4D**). Second, σR4 was displaced toward the β’-ZBD and residues σ^Β^ -H252, E249 and A245 made new contacts with β’-ZBD. The contacts of σR4 with the β’-dock and β-clamp, characteristics of the docked σR4 conformation, were lost (**Figure 4D**). Superposition of the undocked Eσ^B^R4 structure with the published structure of *Mtb* RPc (47, 71) showed that undocked σR4 was incompatible with binding to the -35 element (**Figure 4E)** and should hinder RPc formation on -10/-35 class promoters. Conversely, it should not affect RPc formation on extended -10 class promoters because σR3 positioning was unchanged. Our results revealed the high conformational flexibility of σR4 that may help RNAP to adopt a large spectrum of promoter architectures and provides a target for regulation by σR4-binding transcription factors (48, 72). β-CTT, visible only in the undocked RNAP structure, was missing from all published of *Mycobacterial* RNAP holoenzymes structures, but was observed in the paused transcription elongation complex (TEC) that comprises the elongation factor NusG (60). We hypothesize that β-CTT, found only in a subset of bacterial species, may be implicated in the assembly of the mature RNAP holoenzyme and also in promoter escape when σR4 should be ejected from RNA exit channel.

### Binding of the -10 promoter element to RNAP mimics RbpA-induced σR3-σR4 loading

To investigate the mechanism by which -10 ssDNA induces a conformational change in Eσ^B^, we explored the conformational dynamics of σ^B^ by smFRET. We used a σ^B^ subunit stochastically labeled with the DY547P1 and DY647P1 fluorophores in σR2 (position 151) and σR4 (position 292), as described before(49) (**Figure 5A**). The distances between labels in free (unbound) σ^B^ and σ^B^ in RPo were ∼50 Å (closed σ^B^ conformation, E_PR_ = 0.83, E=0.78) and 83 Å (open σ^B^ conformation, E_PR_ = 0.41, E=0.17), respectively (49) (**Figure 5A**). The distance between the Cα atoms of the σ^B^ residues 151 and 292 in the Eσ^B^/-10 ssDNA complex (docked σ^B^R4), was 68 Å, which matches the distance in the *Mtb* RPo (**Figure 5A**). The distance in the Eσ^B^/-10 ssDNA complex with undocked σ^B^R4, was intermediate (56.6 Å). Previously, we showed that RbpA forms a stable complex with free σ^B^ (42), but does not affect its conformational dynamics (49). Addition of -10 ssDNA to free σ^B^ also had little effect on its conformation (∼70% of molecules remained in the closed state) (**Figure 5B**), in agreement with the cross-linking experiments (**Figure 3E**). Addition of both, -10 ssDNA and RbpA resulted in a multimodal distribution with a peak corresponding to the closed conformation (E_PR_=0.87, 22%), a mean peak at intermediate FRET efficiency (E_PR_=0.63, 44%), and a third peak at low FRET efficiency (E_PR_=0.31, 34%) **(Figure 5B)**. We concluded that cooperative binding of the -10 ssDNA and RbpA to σ^B^ increases its dwell time in a partially open conformation but is not sufficient to stabilize the fully open conformation found in the RNAP holoenzyme. This correlated with the increased affinity of σ^B^ to -10 ssDNA (**Figure 3E-G**). We suggest that binding of -10 ssDNA to the RbpA-σ^B^ complex is stabilized through direct interactions with both proteins. In the holoenzyme (*Mtb* Eσ^B^), σ^B^ was mainly found in its closed conformation as reported earlier (49) (**Figure 5C**). Addition of -10 ssDNA resulted in a significant increase in the low FRET subpopulation (E_PR_=0.31) (**Figure 5C**), leading to a pattern that resembled the one observed in the complex of RNAP with us-fork DNA (49). This suggested that -10 ssDNA binding to domain σ2 stimulates RNAP core-dependent conformational changes in σ^B^ and shifts the equilibrium towards the open σ^B^ conformation with σR4 loaded onto the RNA exit channel. Addition of both -10 ssDNA and RbpA to the *Mtb* Eσ^B^ holoenzyme shifted the equilibrium towards low FRET (E_PR_=0.29) and 20% more molecules adopted the open conformation (**Figure 5C**). This change may reflect an increase in the number of molecules that comprise docked σ^B^R4. We concluded that RbpA, which interacts with -10 ssDNA, β’-ZBD and σ-NCR, fastens the whole complex and stabilizes σ^B^R2 interactions with -10 ssDNA. Stabilization of -10 ssDNA binding hinders the spontaneous collapse of σ^B^ to the closed conformation and favors σ^B^R3-R4 loading to RNAP.

**Figure 5.**
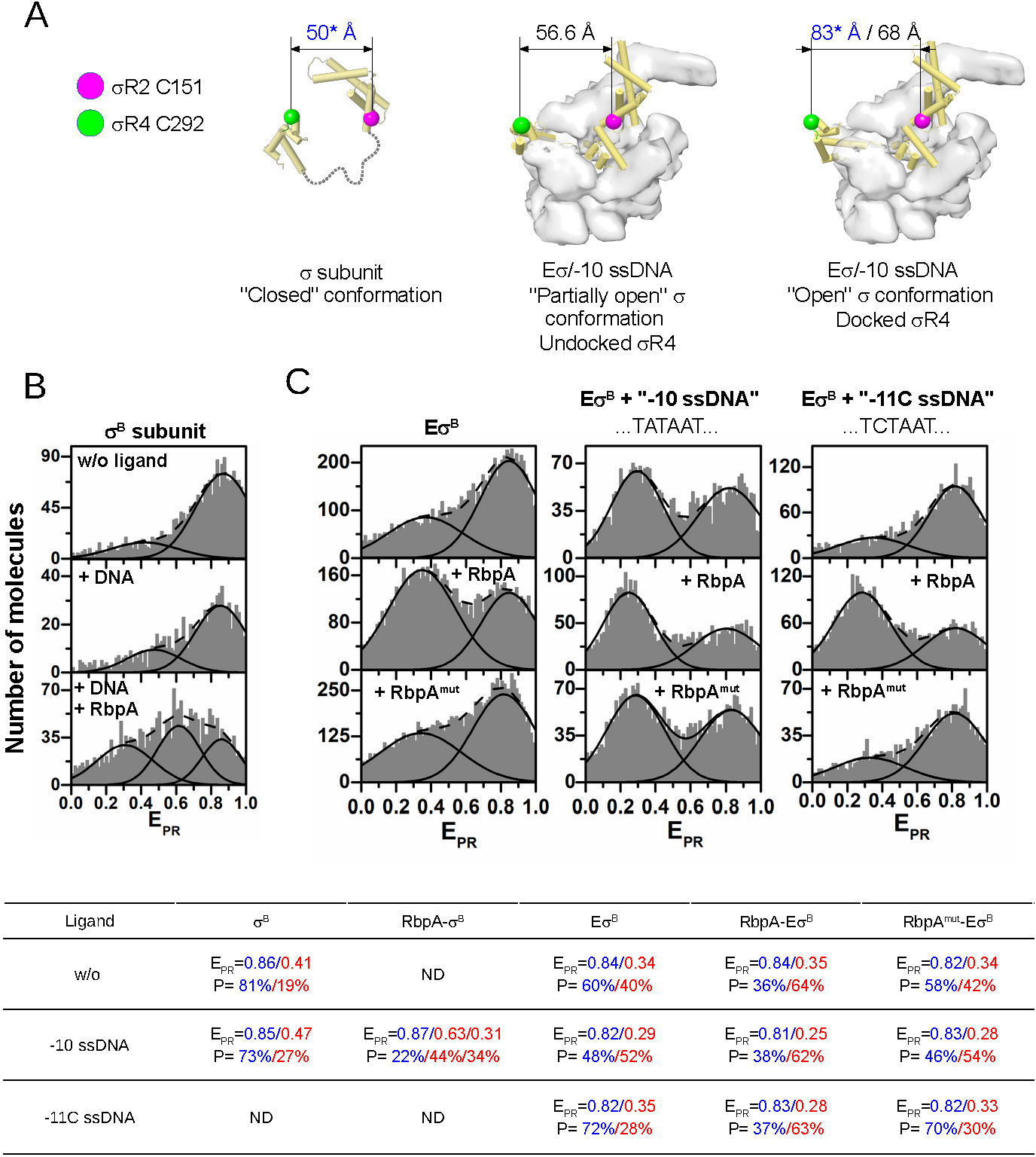
Conformational changes in σ^B^ upon the -10 ssDNA and RbpA binding. (A) Molecular models of the free σ^B^ subunit (ribbons with cylindrical helices) and Eσ^B^ in the undocked and docked conformations. The RNAP core is depicted as a molecular surface in gray. Spheres (pink and green) represent the Cα atoms of the σ^B^ subunit residues labeled with the DY-547P1 and DY-647P1 fluorescent dyes. Distances between dyes, shown above the models, were calculated using the smFRET data (in blue) (49) and from the Eσ^B^ structures reported here (in black). (B) smFRET of free σ^B^ performed without ligands or in the presence of -10 ssDNA (DNA) and RbpA. (C) smFRET of the Eσ^B^ holoenzyme in the presence of -10 ssDNA, -11C ssDNA, wild type RbpA and the RbpA double mutant R88,89A (RbpA^mut^). The table lists the EPR values (E) and percentages (P) of molecules in each histogram peak from panels B and C. ND, not determined.

### Interaction of σ2 with -10 ssDNA and RbpA σ-binding-domain (SID) induces σR3-σR4 loading

To verify that the smFRET efficiency change induced by -10 ssDNA was due to its sequence-specific interaction with σR2, we performed experiments with -11C ssDNA (**Figure 5C**). Addition of -11C ssDNA to the *Mtb* Eσ^B^ holoenzyme did not induce any increase in the low FRET population, suggesting that σ^B^ remained in the closed confirmation. The E_PR_ distribution pattern matched that of the RNAP holoenzyme. Addition of RbpA and of -11C ssDNA to the *Mtb* Eσ^B^ holoenzyme resulted in the increase of the low FRET subpopulation (E_PR_=0.28, 60%), as observed with the RbpA*-*Eσ^B^/-10 ssDNA complex (E_PR_=0.31, 53%) (**Figure 5C**). This is likely due to stabilization of the σ^B^/-10 ssDNA interaction by RbpA that compensates the effect of the -11C mutation. Interaction of RbpA with the σ^B^ subunit NCR (σ^B^-NCR) is essential for transcription activation (46, 47). To verify that the observed effect of RbpA on σ^B^ conformation was due to its interaction with σ^B^-NCR, we performed smFRET experiments with a mutant RbpA harboring R88A and R89A in its SID (**Figure 5C, panels RbpA^mut^**). These alanine substitutions abolish RbpA-σ interactions and RbpA-mediated transcription activation (41). The two mutations completely suppressed σ opening by RbpA. Addition of -10 ssDNA to the RbpA^mut^-Eσ^B^ complex produced the same pattern as -10 ssDNA alone, showing the stimulating effect of RbpA loss. Finally, combining mutant -11C ssDNA with RbpA^mut^ resulted in a E_PR_ distribution similar to that of the Eσ^B^ holoenzyme. Based on these results, we concluded that σ loading is induced by the specific interaction of RbpA and -10 ssDNA with their respective binding sites on the RNAP holoenzyme and that RbpA and -10 ssDNA are interchangeable (**Figure 5C bottom**).

### The β subunit flap is essential for RbpA-driven σR3-σR4 loading

Binding of σR4.2 to the β subunit flap-tip-helix (β-FT) domain (**Figure 6A**) is essential for positioning σR4.2 to interact with the -35 sequence element (73), but not for the *Mtb* Eσ^B^ immature holoenzyme formation (50). To determine whether the β-FT/σR4.2 contact was implicated in RbpA and -10 ssDNA-mediated σR3-σR4 loading, we performed transcription assays and smFRET measurements with the mutant *Mtb* E^ΔFT^ in which the β subunit residues 811-825 were deleted. The mutant *Mtb* E^ΔFT^ core is active in the promoter-independent initial transcription assays with a DNA scaffold template (61). Here, we first tested the activity of the *Mtb* E^ΔFT^σ^B^ holoenzyme in transcription runoff assays with the RbpA-dependent *sigA*P promoter that contains an almost perfect -10 sequence element (5 of 6 matches) and displays weak homology to the -35 sequence element (3 of 6 matches) (**Figure 6B,C**) (43). The β-FT deletion almost fully suppressed the transcription stimulation by RbpA, suggesting that RbpA functioning strongly depends on the β-flap. Previous studies on *Ecol* RNAP showed that β-FT is dispensable for transcription initiation from the extended -10 class promoters without the -35 motif. In this context, interaction of σR3 with the T_-17_R_-16_T_-15_G_-14_ motif of the extended -10 element is essential for recognition of the extended -10 class promoters, while interaction of σ4.2 with the -35 motif is dispensable (74). The extended -10 motif stimulates transcription initiation by Eσ^B^ independently of RbpA (45). When we tested the activity of the mutant *Mtb* E^ΔFT^σ^B^ holoenzyme on a synthetic *sigA*P promoter variant that contained the T_-17_R_-16_T_-15_G_-14_ motif (*sigA*P-TGTG, **Figure 6B,C**), we found that the β-FT of *Mtb* RNAP was essential for transcription from the *sigA*P-TGTG promoter (**Figure 6B,C**), unlike what observed for *Eco* RNAP (73). Addition of RbpA restored the *Mtb* E^ΔFT^σ^B^ holoenzyme activity to the level of wild type RNAP in the absence of RbpA. Our results indicate that despite the high level of conservation in the core RNAP structure and function, the *Eco* paradigm might not be applicable to other bacterial species. The effect of the β-FT deletion was reminiscent of the effect of the deletion of its partner σR4.2 that leads to transcription inhibition from *sigA*P-TGTG (45).

**Figure 6.**
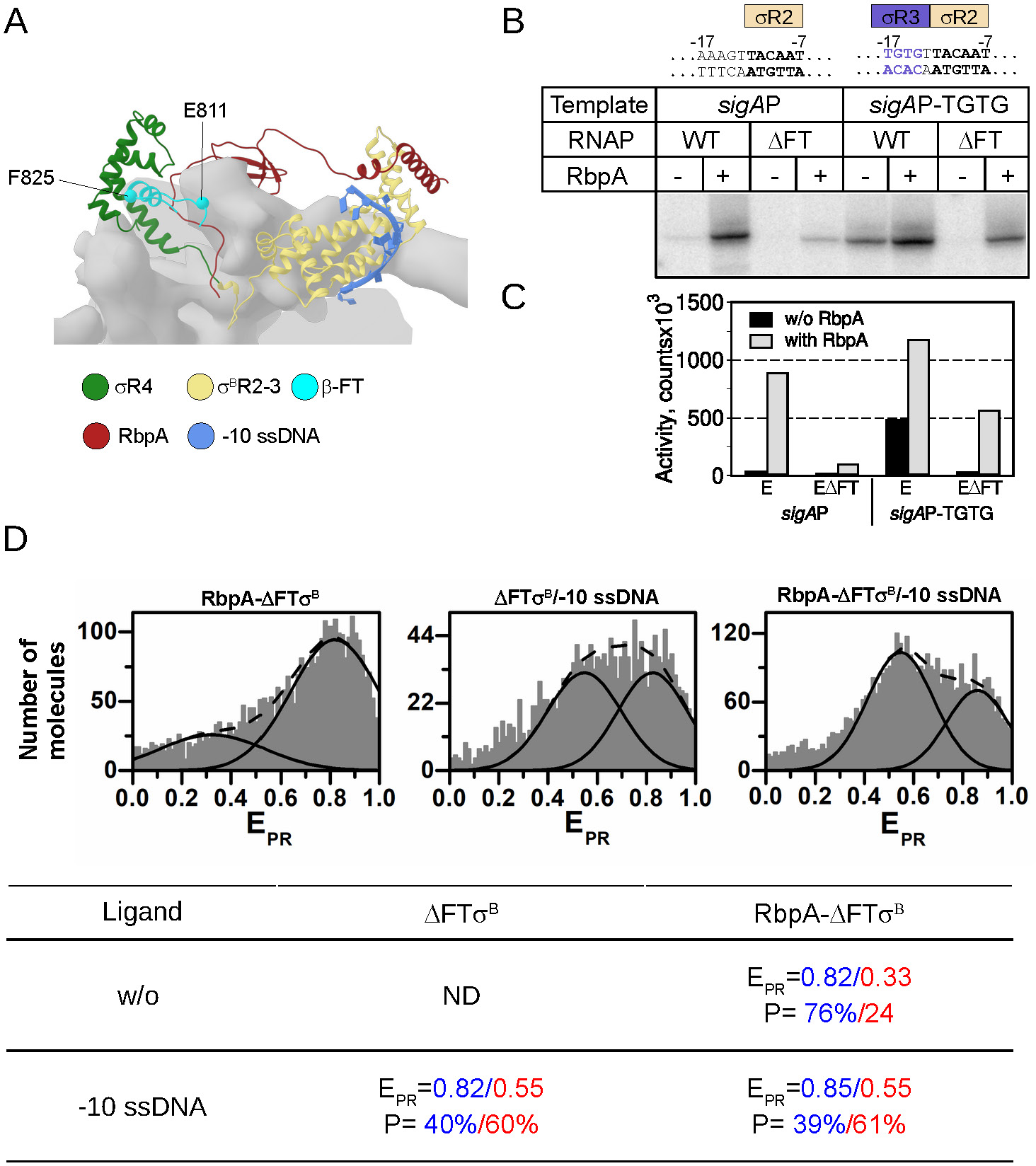
The RNAP β-flap is essential for σ loading induced by RbpA and by -10 ssDNA. (A) Cartoon showing the σ^B^ 4/β-FT interactions. Molecular model of Eσ^B^/-10 ssDNA in complex with RbpA. The RNAP core is depicted as molecular surface in gray. The σ^B^ subunit, RbpA and β-FT are shown as ribbons models. (B) Effect of β-FT deletion on run-off transcription from the *sigA*P and extended -10 type *sigA*P-TGTG promoters. The difference in promoter sequences is shown on the top. The σ^B^ subunit regions that interact with promoter motifs are depicted as rectangles above the DNA sequences. (C) Bar graph showing the quantification of the run-off RNA products in panel B. (D) smFRET analysis of the mutant *Mtb* E^ΔFT^σ^B^ holoenzyme (lacking the RNAP β-FT) in the presence of -10 ssDNA and RbpA. The table lists the EPR values (E) and percentages (P) of molecules in each peak in the panel D. ND, not determined.

In agreement with the transcription assay findings, smFRET experiments showed that binding of RbpA to the mutant *Mtb* E^ΔFT^σ^B^ holoenzyme had little effect on E_PR_ distribution (**Figure 6D**), compared with RbpA binding to wild type *Mtb* Eσ^B^ (**Figure 5C)**. Only a minor subpopulation of molecules (24%, vs 64% for wild type) were in the low FRET state, corresponding to the open σ^B^ conformation (E_PR_ = 0.34). The distribution of FRET states was similar to that of the *Mtb* Eσ^B^ holoenzyme in the absence of RbpA (**Figure 5C**). We concluded that in the *Mtb* E^ΔFT^σ^B^ holoenzyme, most σ^B^ molecules remained in the closed conformation. Therefore, when σ^B^R4.2/β-FT interaction is disrupted, RbpA cannot induce the σ^B^ conformational change essential for RPo formation. These results perfectly fit with the mechanism where RbpA promotes the formation of the contacts between σR4.2 and β-FT that are essential for the correct σR3 positioning relative to the extended -10 motif.

### The β subunit flap is essential for -10 ssDNA driven σR3-σR4 loading

Then, we tested the effect of β-FT deletion on -10 ssDNA-induced loading of σR3-σR4 to RNAP. Addition of the -10 ssDNA to *Mtb* E^ΔFT^σ^B^ resulted in a broad E_PR_ distribution of the molecules that could be fitted with two Gaussian models. It indicated an equilibrium between a σ^B^ intermediate in a partially opened conformation (E_PR_=0.55; 52%) and the closed confirmation (E_PR_=0.82; 48%). We observed no peak with E_PR_ =0.31, a feature of the open σ conformation (see **Figure 6D, panel “1′FTσ^B^/-10 ssDNA”**). Thus, similar to what observed for RbpA, β-FT is required for the -10 ssDNA-induced conformational change in the σ subunit. Upon addition of both RbpA and -10 ssDNA (**Figure 6D, panel “RbpA-1′FTσ^B^/-10 ssDNA”**), the equilibrium was further shifted towards the intermediate σ^B^ conformation (E_PR_=0.54, 63%). These results suggest that both RbpA and -10 ssDNA bind to the mutant *Mtb* E^ΔFT^σ^B^, but cannot promote the correct positioning of σR4 due to the absence of β-FT, its anchoring point. Furthermore, RbpA and -10 ssDNA remodel the σ subunit through a similar mechanism by targeting σ2. The effect of -10 ssDNA on σ conformation in the absence of the σR4/β-FT contact on the RNAP core suggests that σ2 is exposed for binding to ssDNA. We propose that ssDNA binding to σ2 destabilizes the overall fold of the σ subunit, which allows regions σ3 and σ4 to interact with their respective binding sites on the RNAP core.

### Clamp movements and σR4 swinging do not correlate

To determine whether there is a correlation between σR4 docking and clamp closure/swiveling, we performed pairwise cluster analysis (nine clusters) on variability components generated by 3DVA with the full consensus II map (**Supplementary Figure S6**). We compared the variability components 1 (C1, σR4 swinging) with the variability components 0 (C0, clamp swiveling) and 2 (C2, clamp closing) and also C0 with C2. For each component pair, we sorted particles in nine clusters (**Figure 7A-C, Supplementary Figure S6C**). We used the resulting series of cryo-EM maps for rigid body refinement of the consensus I molecular model. Clamp rotation angle measurement has been widely used to characterize RPo conformational dynamics (30, 34, 75). However, results depend on the choice of reference structure. Here, to characterize RNAP domain movements, we measured the distances between reference points in each of the refined models that provide absolute characteristics for a given structure independently of the reference choice (**Supplementary data table 1)**. To quantify the σR4 domain movement, we measured the distances between the Cα atoms of σ-I247 in σR4 and β’-I469 in β’ dock. To quantify clamp/lobe movements, we measured the distances between β-G284 in the β lobe and β’-K123 in the β’ clamp head and β’-L360 in the clamp core (**Figure 7D**). By plotting the distance ranges between σ-I247 and β’-I469 against the distances between β-G284 and clamp (β’-K123, β’-L360) (**Figure 7A,B, Supplementary data table 1**), we found that σR4 adopted two distinct conformations with average distances of ∼7 Å (docked σR4) and ∼17 Å (undocked σR4). We did not detect any intermediate state. This suggests that such states are short-lived. Moreover, we did not find any correlation between clamp and σR4 movements. Unlike σR4, clamp movements were gradual and displayed a clear correlation between swiveling and opening/closing (**Figure 7C, Supplementary data table 1**): the open clamp state adopted a more swiveled conformation and the closed clamp state an unswiveled conformation. We concluded that after σR3 binding to the β-lobe and σR4 insertion into the RNA exit channel, clamp adopts preferentially an unswiveled/closed conformation, but remains dynamic. This is essential for DNA entry into the active site cleft during the isomerization from RPc to RPo.

**Figure 7.**
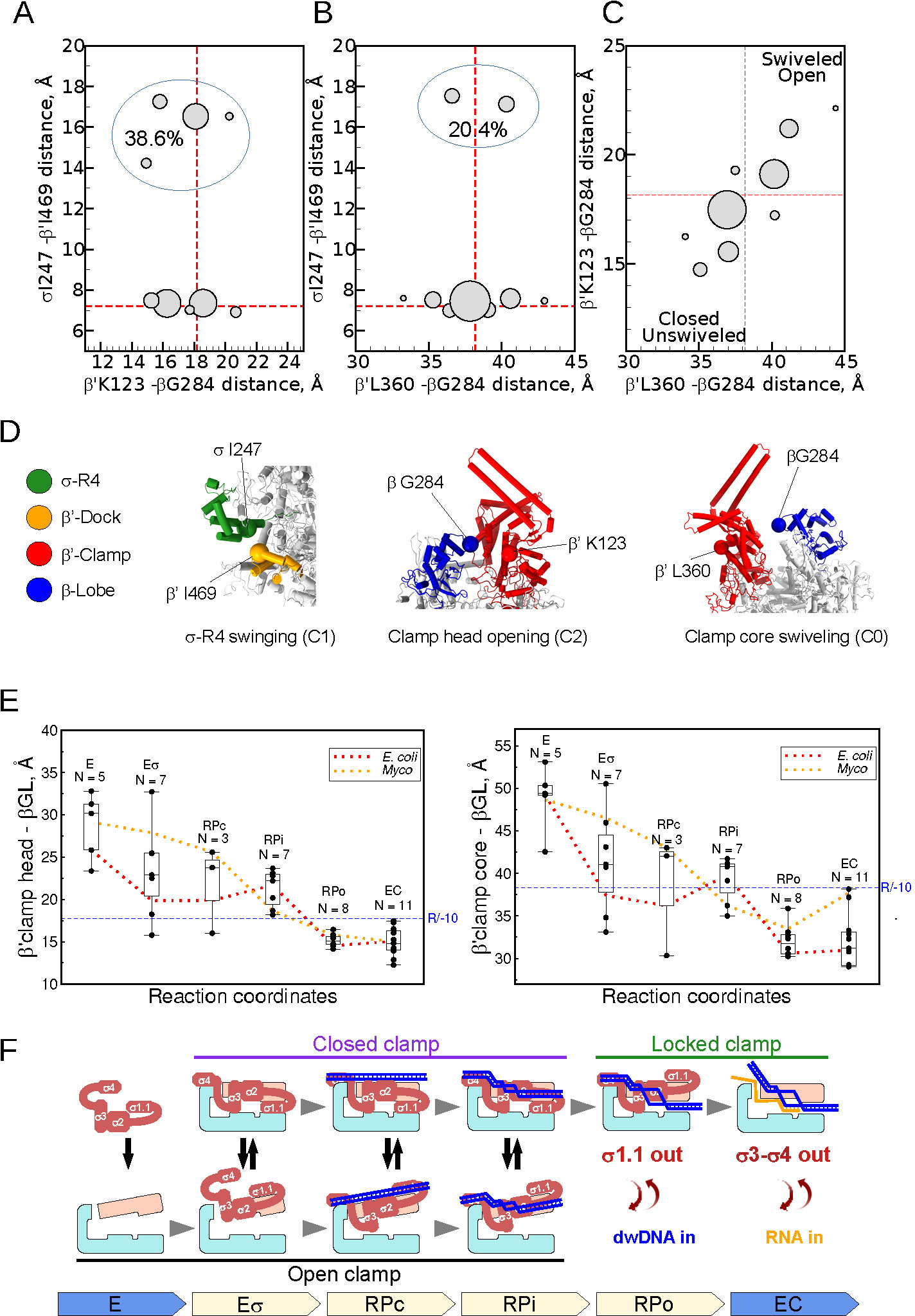
Correlation between σ loading, σR4 swinging and clamp motions. (A,B,C) Graphs showing the distances between σR4 and β’-dock and β’CH and β-lobe in the nine clusters produced in the 3DVA. Diameter of the circles is proportional to the cluster size (D) Molecular models showing the above mentioned regions; the Cα atoms of the residues used for the distance calculations are shown as spheres. (E) Box plots showing the compilation of the clamp-lobe distances from published cryo-EM structures of RNAP plotted in function of the transcription cycle progression (Reaction coordinates). Dotted lines show the mean values for *E. coli* and *M. tuberculosis* (*Myco*), respectively. N, number of data points (N) used for each reaction step. (F) Model depicting the changes in RNAP clamp (light salmon) and σ subunit (Indian red) conformations throughout the transcription cycle. Promoter DNA is depicted as a blue ladder, RNA as an orange line. Each complex type (E; Eσ, RPc, RPi, RPo, EC) and the reaction direction are indicated by arrows at the bottom.

### Clamp dynamics gradually decrease during progression of the transcription cycle

To relate the observed β‘-clamp/β-lobe movements in the RNAP/-10 ssDNA complex with RNAP conformational dynamics throughout the transcription cycle, we analyzed the β’-clamp/β-lobe states in forty published cryo-EM structures of bacterial RNAPs, alone and in complex with various transcription factors (**Figure 7E, supplementary data table 2**). The ranges of distances from the β lobe (β-G284) to the β’ clamp head (β’-K123) and to the clamp core (β’-L360) are presented as box plots for each transcription cycle step: assembly of the core (E) and σ subunit into the RNAP holoenzyme (Eσ), promoter complex formation (RPc, RPi, RPo), and elongation (EC). The distance analysis showed that the RNAP core adopted the largest open clamp state with a wide range of distances (amplitude ∼10 Å). After RNAP holoenzyme assembly, the clamp displayed a broader range of states (amplitude ∼17 Å), from wide open (matching the RNAP core state) to closed (matching the RPo state). The immature *Mtb* Eσ^B^ holoenzyme, in which domains σR3-σR4 are unbound from their respective sites in the core, adopted a wide open clamp state (distances of 32.7/50.6 Å), similar to the RNAP core (50, 76). We concluded that binding of the σR3 domain to the RNAP β-protrusion and insertion of the σR4 domain in the RNA channel increase the conformational mobility of the core enzyme and promote clamp closure. RNAP binding to promoter DNA duplexes restrained clamp movements in RPc while keeping the wide range of states characteristics of holoenzymes. Nucleation of promoter DNA melting in RPi led to a decrease in clamp mobility (amplitude ∼5 Å); however, the clamp could still adopt partially open states. Upon RPo formation, replacement of the σR1.1 in the DNA channel by the downstream promoter DNA duplex froze the clamp in the closed state and limited its conformational dynamics (amplitude ∼2 Å) (**Figure 7E,F**). After the promoter escape to elongation, σR3-σR4 are replaced by nascent RNA that favors the σ subunit dissociation. Overall, clamp dynamics increased, but the clamp remained mainly closed over DNA template. We observed the most closed clamp state in *Eco* his paused TEC and the most unswiveled clamp state in *Eco* elemental paused TEC. Although the general tendency was towards a decrease in clamp-lobe distances over the reaction coordinates, we observed some differences in distance distribution in every dataset due to uncoupling between clamp closing and clamp swiveling (most noticeable for RPo and EC). Plotting separately the average distances of the *Eco* (red dotted line) and *Mtb* (orange dotted line) datasets (**Figure 7E**) highlighted a large data divergence particularly for the RNAP core (E) and RPc. Yet, due to the small number of available structures, this difference was not significant. The clamp-lobe distance range of the *Mtb* Eσ^B^/-10 ssDNA complex (17.4/38.6 Å, amplitude of ∼5 Å) positioned it between RPi and RPo (**Figure 7E**, blue dashed line, R/-10). We suggest that RNAP conformational state in this complex may correspond to that in the transient intermediate on the path to RPo. Unlike in RPi and the *Mtb* Eσ^B^/-10 ssDNA complex, in RPo, region σR1.1 is displaced by dwDNA and the downstream part of the transcription bubble (dw-fork DNA) that enters the DNA channel. Thus, we concluded that binding of dw-fork DNA locks the clamp in the closed state, which is retained during elongation (EC) (**Figure 7F**). The results of our analysis are in perfect agreement with the smFRET data showing a wide open clamp in the RNAP core, a partially closed clamp in the RNAP holoenzyme, and a fully closed/locked clamp in RPo (26).

## DISCUSSION

The principal finding of our work is that the σ subunit functions as a sensor of regulatory signals from the ssDNA effector, leading to isomerization between the open and closed RNAP clamp conformations . Therefore, as proposed by Ishihama (77), several decades ago, σ is not just a promoter-recognition-melting factor, but also a major regulator of RNAP conformational dynamics. We suggest that in solution, the default “relaxed” conformations of the clamp in the RNAP core and of the σ subunit are open/swiveled and closed with masked DNA binding regions, respectively. Bringing together the two partners leads to σ opening and RNAP clamp closing/unswiveling and consequently to the formation of a “stressed” RNAP, competent to initiate transcription on promoter DNA.

### The σ-mediated maturation of the RNAP active site and analogy with NusG/RfaH

The swiveled clamp state was first described as an attribute of the PECs (58) regulated by the universal transcription factor NusG/Spt5 and its paralog RfaH (59, 60, 78). The anti-pausing *Eco* NusG suppresses swiveling by contacting β-GL (59). We showed that σ subunit domains loading onto the nucleic acid binding channels of RNAP induced a catalytically proficient unswiveled conformation of the active site. This finding provides a structural rational for the previously reported anti-pausing activity of *Eco* σ^70^ and *Mtb* σ^B^ during initial transcription (61) and reveals a remarkable functional similarity between σ factors and elongation factors that act via universal RNAP domains: β’-CH, protrusion, β-flap and β-GL.

### Linking σ loading and clamp closure

The clamp closure/unswiveling observed in the Eσ^B^/-10ssDNA complex can be induced by loading of σ domains or/and by -10 ssDNA binding. As -10 ssDNA does not make stable contacts with the RNAP core, we propose that loading of σR3-σR4, and probably of σ1.1, promotes clamp closure. This conclusion is supported by the comparison of the published cryo-EM structures of RNAPs. (1) The clamp is open in the *Mtb* RNAP core and RNAP holoenzyme that comprises σ subunits with unloaded σR3-σR4. (2) Similarly, in the published structures of *Mycobacterium smegmatis* RNAP core and holoenzyme with unloaded σ^A^R4, the clamp is in the open conformation. (3) In the RbpA-Eσ^A^ complex, which contains fully loaded σR3-σR4, the clamp adopts a more closed conformation than in the RNAP core. (4) In *Eco*, the clamp is open in the RNAP core and closed in the Eσ^70^ holoenzyme that contains fully loaded σR3-σR4. Yet, the smFRET findings suggested that the clamp adopts mainly an open state in both *Eco* RNAP core and holoenzyme, unlike the closed state in RPo. Conversely, the cryo-EM analysis of *Eco* Eσ^70^ showed the clamp in a closed state, as found in RPo. In addition, smFRET showed a reduction of the opening-closing dynamics of σ^70^ in RPo compared with the holoenzyme (49). This discrepancy might reflects the fact that the published structures represent a “consensus” conformation of RNAP derived from a conformational ensemble. The clamp high conformational mobility, modulated by the transcription factor TraR, has been observed in the *Ecol* Eσ^70^ holoenzyme (75). We interpret these findings as the evidence of a larger amplitude of clamp motions in the holoenzyme induced by σ loading. In *Mtb* RNAP, the equilibrium between clamp states is shifted towards the open conformation due to the weak binding of σR3-σR4. In *Ecol* RNAP, the equilibrium is shifted towards the closed conformation due to the stable binding of σR3-σR4.

### Stabilization of the closed clamp state by the σ/ssDNA interaction and path to RPo

The presence of a DNA template drastically reduces the clamp and σ domain conformational mobility in RPi/RPo (23, 24). In the Eσ^B^/-10 ssDNA complex, the clamp adopted a more closed state than in the *Mtb* RNAP holoenzymes, but more open than in RPo. We infer that ssDNA binding to RNAP restricts the clamp dynamics and shifts its conformational equilibrium towards a closed state, similar to that observed in RPi. Indeed, all early intermediates (RPi) with the melted -10 element show a partially closed clamp state (10, 34, 64). In RPo, after the replacement of σR1.1 by dwDNA, the clamp becomes locked in the closed state and remains locked during elongation (26) (**Figure 7F**). This scheme fits well with the *Mtb* model, but may differ from that of *Eco* (**Figure 7E, dotted lines**). Moreover, the conformational equilibrium between clamp states can be influenced by the experimental conditions, such as type of promoter DNA template, divalent ion concentration and presence of transcription factors (e.g. *Eco* TraR) (26, 75).

### Difference in RNAP holoenzyme assembly pathway in *E. coli* and *M. tuberculosis*

Despite decades of studies, the mechanism of RNAP holoenzyme assembly remains poorly understood, and the exact order of events of the σ subunit loading to the RNAP core is unknown (77, 79, 80). Our results suggest that assembly starts after σR2 binding to β’-CH on the clamp and σR1.1 insertion into the dwDNA channel. The open clamp conformation of the RNAP core and β-flap wobbling allow the entry of the C-terminal domains of σ into RNAP main cleft. Binding of σR3 to the β-protrusion ties the RNAP pincers together and favors clamp closure. Finally, σR4 inserts into the RNA exit channel and binds to the β-flap and β-clamp, stabilizing the whole system. On the basis of the high conservation of the core binding regions of group I/II σ subunits and their respective binding sites on RNAP, the assembly pathway should be universal for all bacteria, but may differ in the nature of bottleneck steps delimited by lineage-specific insertions/deletions in σ and RNAP. For example, in *Eco,* σ^70^ loading onto the RNAP core is a spontaneous process that does not require any additional co-factor. Conversely, in *Mtb,* σ^A^ and σ^B^ loading onto the RNAP core is RbpA-dependent. This RbpA dependency is a feature of the *Mtb* RNAP core and not of the σ subunit because assembly of a chimeric holoenzyme (*Mtb* σ^B^ and *Eco* RNAP core) proceeds without RbpA (49).

### Allosteric switch in σ and stimulation of binding to the non-template DNA strand binding by RNAP core

A large number of biochemical studies on *Eco* and *Taq* models reported a core-induced conformational change in the group I σ subunit leading to unmasking of its DNA binding regions and to stimulation of -10 ssDNA binding (8, 66, 81). Our smFRET and structural studies in *Mtb* and *Eco* suggest that the only detectable conformation change in σ is the movement of σR3-σR4 away from σR2, providing a time window for ssDNA binding. The RNAP core stabilizes -10 ssDNA/σ2-interactions by favoring contacts between DIS and σR1.2, thus preventing σ closing. In *Mtb*, RbpA tethers -10ssDNA to σR2 and prevents σ closing. Similarly, it has been shown that a DNA aptamer containing -10 ssDNA induces *Taq* σ^A^ opening (82). However, a σ^70^ fragment that lack regions σR3-σR4 (σ2) displayed enhanced binding to -10 ssDNA in the presence of the RNAP core (14) or the β’ clamp fragment alone (18). Our analysis of cryo-EM structures suggest that the RNAP core alone does not induce any global conformational change in σR2. Conversely, -10 ssDNA induces W-dyad isomerization that may stabilize contacts between DIS and σR1.2. Although the overall σ2 fold is conserved in all group I-II σ subunits, the regulation of -10 element binding may vary among species due to lineage-specific insertions in σ-NCR. For instance, insertions in the *Eco* σ^70^ and *Taq* σ^A^ σ-NCR would clash with β’-CH and impose a conformational change in σ2 upon holoenzyme assembly. A plausible hypothesis is that σR1.2 N-terminal α-helix (*Mtb* σ^B^ residues 26-36), which is attached to the main body of σ2 through an unstructured linker, may be displaced in free σ and will not form optimal contacts with DIS. Binding to the RNAP core may lock the σR1.2 α-helix in the correct conformation that is optimal for DIS binding.

### Multiple RbpA roles in transcription initiation

RbpA stabilizes the RNAP holoenzyme (43), stimulates promoter DNA melting (42, 43) and slows down promoter escape (47, 83). Our results explain the multiple roles of RbpA in transcription initiation and suggest that RbpA acts from RNAP assembly to promoter escape by stabilizing the σ2/-10 element, σ3/extended -10 element and σ4/-35 element interactions. Specifically, RbpA induces the optimal fit of RNAP to -10/-35 elements upon RPc formation. Then, RbpA-mediated anchoring of σR2 to -10 ssDNA should stimulate formation of the transcription bubble during isomerization from RPc to RPo. The latter implies that RbpA should slow down promoter escape by inhibiting the disruption of σR2/-10 contacts, in agreement with the kinetics studies (83) .

## DATA AVAILABILITY

The data underlying this article will be shared on reasonable request to the corresponding author. The cryo-EM density maps and model coordinates reported in this article are available in the Electron Microscopy Data Bank (EMDB) and Protein Data Bank (PDB) and can be accessed with the accession codes: EMD-50508 (Consensus-I map); 9FJP (Consensus-I model); EMD-50509 (Consensus-II map); EMD-50510 (σR4-docked state map); 9FJR (σR4-docked state model); EMD-50511(σR4-undocked state map); 9FJS (σR4-undocked state model); EMD-50512 (swiveled clamp state map); EMD-50514 (unswiveled clamp state map).

**Supplementary Data are available at NAR online**

## AUTHOR CONTRIBUTIONS

Conceptualization, K.B.; Data Curation, K.B., E.M. and R.K.V; Formal Analysis, K.B., E.M., R.K.V., N.M, and M.B.; Investigation, R.K.V., N.M. Z.M., and K.B.; Methodology, K.B., E.M. and N.M.; Supervision K.B. and EM.; Visualization K.B. and R.K.V.; Validation, K.B., E.M., R.K.V. and M.B.; Writing – original draft, K.B.; Writing – review & editing, K.B., E.M. R.K.V., N.M. and M.B.; Funding Acquisition, K.B. and E.M.

## ACKNOWLEDGMENTS

We thank Achillefs Kapanidis and Abhishek Mazumder from Oxford University for critical reading of the manuscript and helpful suggestions. We thank Franck Godiard for the collection of negative stained EM data at the MEA platform, University of Montpellier. Cryo-EM data were collected at the Center for Integrative Biology, IGBMC, Strasbourg. The smFRET data were collected at the PIBBS facility, member of the France-BioImaging infrastructure.

