## Supplementary material for "Single-stranded DNA drives σ subunit loading onto RNA polymerase to unlock initiation-competent conformations": Supplemantal Tables and Figures

**Supplementary Table S1. DNA oligonucleotides**

| Oligo name | Sequence | Label | Experiment |
|---|---|---|---|
| -10 ssDNA | 5’-TGCGTATAATGTGTGGA-3’ | - | Cryo-EM, smFRET |
| -10 ssDNA | 5’-[Cyanin5]TGCGTATAATGTGTGGA-3’ | Cy5 | DNA-protein cross-linking |
| -10 ssDNA | 5’-[6FAM]TGCGTATAATGTGTGGA-3’ | Fluorescein | MDS |
| -10 ssDNA truncated | 5’-TATAATGTGTGGA[Cyanin5]-3’ | Cy5 | DNA-protein cross-linking |
| -11C ssDNA | 5’-TGCGTCTAATGTGTGGA-3’ | - | smFRET |
| -11C ssDNA | 5’-[6FAM]TGCGTCTAATGTGTGGA-3’ | Fluorescein | MDS |
| *SigA*P us-fork top strand | 5'-CGCTCGGGCTGTACTCGTGCGCAGTAAAGTTACAATGGTC-3’ | - | EM |
| *SigA*P us-fork bottom strand | 5'-AACTTTACTGCGCACGAGTACAGCCCGAGCG-3’ | - | EM |
| RbpAR88R89A forward primer | 5'-GGGACATGCTGCTGGAGGCCGCTTCCATCGAAGAACTCG-3' | - | site-directed mutagenesis of *rv2050* |
| RbpAR88R89A Reverse primer | 5'-CGAGTTCTTCGATGGAAGCGGCCTCCAGCAGCATGTCCC-3 | - | site-directed mutagenesis of *rv2050* |

**Supplementary Table S2 cryo-EM data collection, refinement and validation statistics**

| | **Consensus-I** | **Consensus-II** | **σR4-docked** | **σR4-undocked** | **Clamp swiveled** | **Clamp unswiveled** |
|---|---|---|---|---|---|---|
| **Data collection** | | | | | | |
| Pixel size (Å) | 0.862 | | | | | |
| Voltage (kV) | 300 | | | | | |
| Electron dose ($e^{-1}$ Å $^{-2}$) | 55.735 | | | | | |
| Defocus range (μm) | -0.8 - -2.5 | | | | | |
| **Reconstruction** | | | | | | |
| Particles used in reconstruction | 290,345 | 167,825 | 72,799 | 67,957 | 21,873 | 36,319 |
| Map resolution (Å) FSC threshold 0.143 | 3.19 | 3.33 | 3.43 | 3.48 | 4.33 | 3.79 |
| **Refinement** | | | | | | |
| Resolution FSC threshold 0.5 | 3.4 | - | 3.7 | 3.7 | - | - |
| Map CC (whole map) (volume) | 0.81 | - | 0.82 | 0.79 | - | - |
| Map CC (peaks) | 0.74 | - | 0.79 | 0.76 | - | - |
| **RMSD** | | | | | | |
| Bond length (Å) | 0.004 | - | 0.004 | 0.004 | - | - |
| Bond angle ( ° ) | 0.931 | - | 0.959 | 0.929 | - | - |
| **Ramachandran Plot** | | | | | | |
| Preferred regions (%) | 97.23 | - | 97.08 | 97.02 | - | - |
| Allowed regions (%) | 2.77 | - | 2.92 | 2.98 | - | - |
| Outliers (%) | 0.00 | - | 0.00 | 0.00 | - | - |
| **Validation** | | | | | | |
| MolProbity score | 1.21 | - | 1.47 | 1.38 | - | - |
| All-atom clashscore | 2.8 | - | 5.67 | 4.31 | - | - |
| Rotamer outliers (%) | 0.74 | - | 0.18 | 0.95 | - | - |
| EM accession | EMD-50508 | EMD-50509 | EMD-50510 | EMD-50511 | EMD-50512 | EMD-50514 |
| PDB accession | 9FJP | - | 9FJR | 9FJS | - | - |

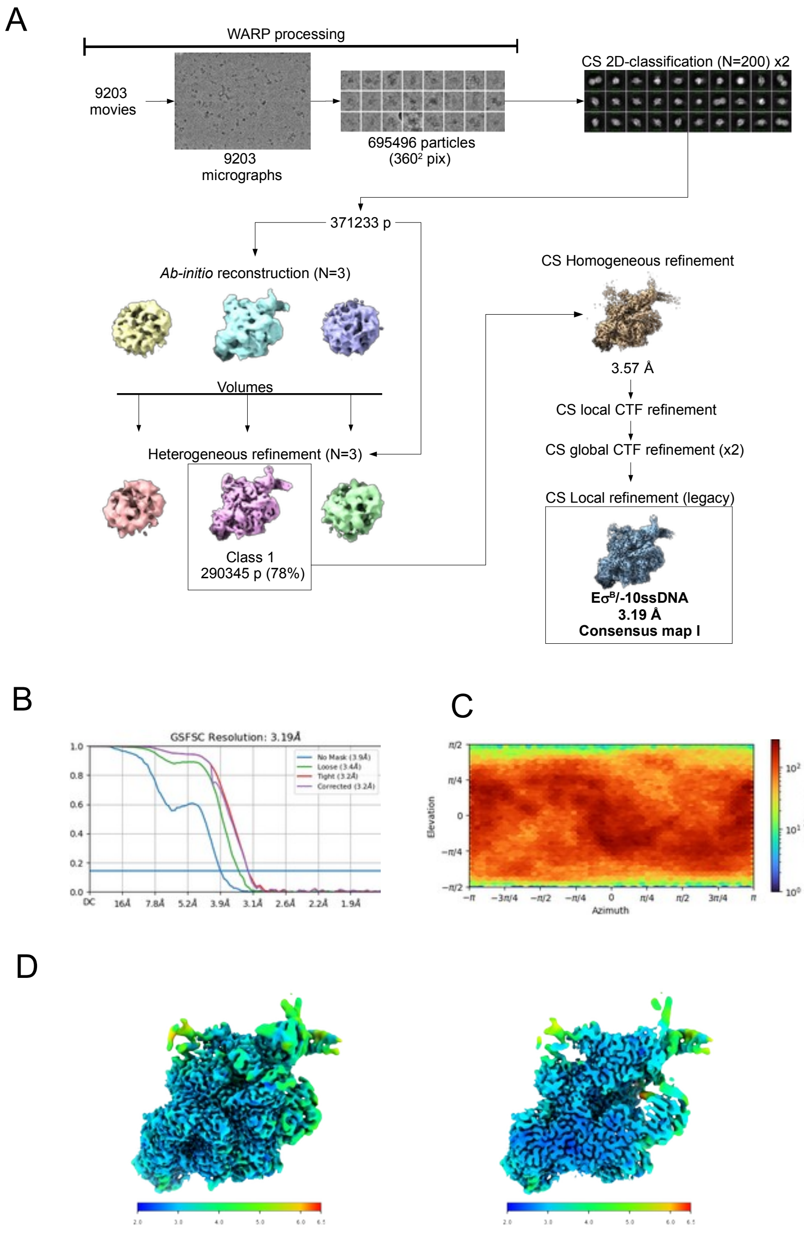


**Figure S1. Reconstruction and characterization of the consensus-I map.**
(A) CryoSPARC pipeline for consensus-I map. (B) Gold-standard FSC calculated for the map in cryoSPARC v3.3.2. The dotted line shows the 0.143 FSC cutoff. (C) Angular distributions for particles projections calculated in cryoSPARC and presented as a heat map. (D) Cryo-EM density map and sliced map (on the right) colored according to the local resolution calculated at 0.143 FSC.

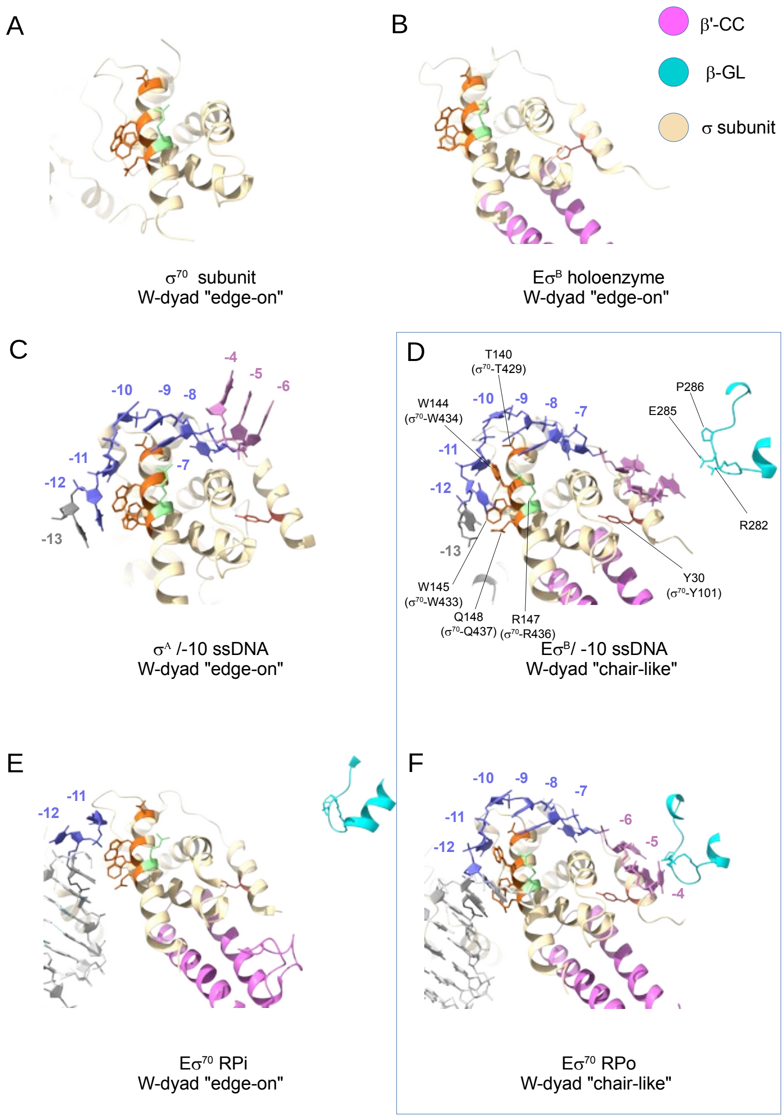


**Figure S2 Architecture of the σ subunit ssDNA-binding interface at different steps of transcription initiation.**

(A) Structure of the *E. coli* $\sigma^{70}$ fragment (PDB:1SIG). (B) Structure of the Mtb $E\sigma^{B}$ holoenzyme (PDB:7PP4). (C) Structure of the *T. aquaticus* $\sigma^{A}$/-10 ssDNA complex (PDB: 3UGP). (D) Structure of the E $\sigma^{B}$/-10 ssDNA complex (current study). (E) Structure of the *E. coli* $E\sigma^{70}$ RPi (PDB: 6PSR) (F) Structure of the *E. coli* $E\sigma^{70}$ RPo (PDB: 7MKD). Ribbon models of the σ subunit colored wheat, β' subunit clamp helices (β'-CH) in pink, β subunit gate loop (β-GL) in cyan. The key residues of σ implicated in -10 recognition and isomerization of RPc to RPo are shows as stick molecular model colored in orange: W-dyad, Q148 ($\sigma^{70}$-Q437, (Waldburger *et al*, 1990), T140 ($\sigma^{70}$-T429) critical for promoter melting (Schroeder *et al*, 2008; Waldburger & Susskind, 1994) and green R147 ($\sigma^{70}$-R436, (Fenton *et al*, 2000). Y30 ($\sigma^{70}$-Y101) is essential for stimulation of the -10 binding by the RNAP core (Zenkin *et al*, 2007). The conserved residues of β-GL implicated in RPo formation are show as stick molecular model.

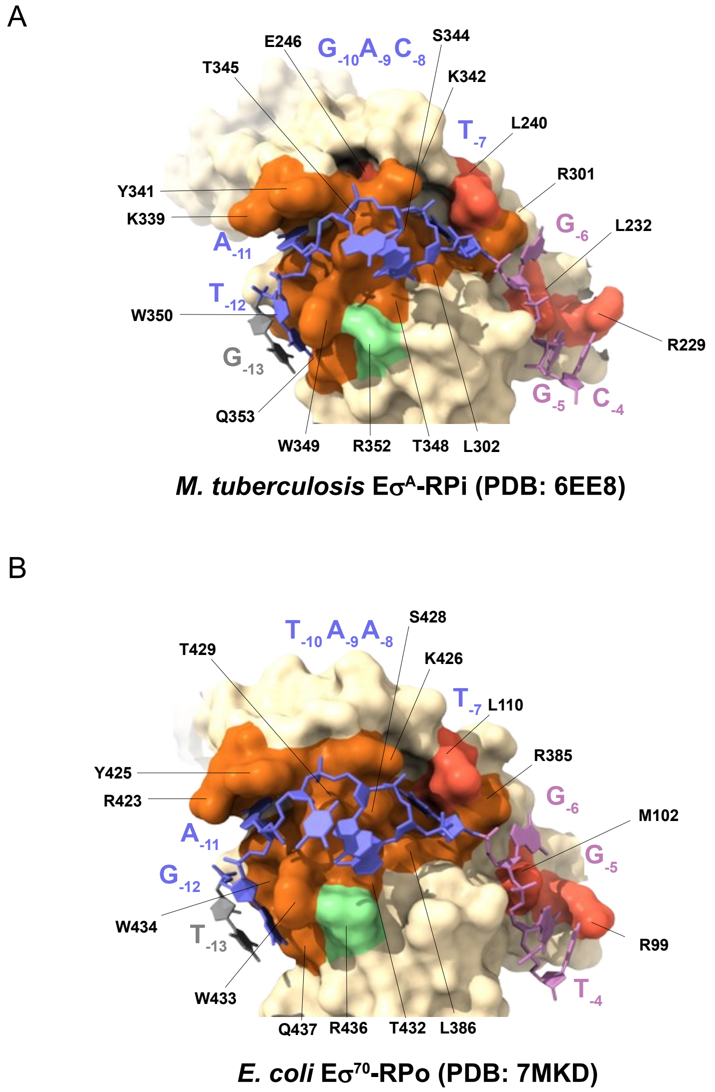


**Figure S3. Architecture of the σ/-10 ssDNA interactions in *Mtb* RPi and *E. coli* RPo**

Comparison of the of the σ/ssDNA binding interfaces in RPo formed by *Mtb*E$\sigma^{A}$ (panel A, PDB: 8EE8) and *E. coli* E$\sigma^{70}$ (panel B, PDB: 7MKD). The σ domain 2 is shown as molecular surface colored in wheat. Residues interacting with ssDNA are colored in orange (σ region 2) and tomato (σ region 1.2). $\sigma^{A}$-R352 and $\sigma^{70}$-R436 making holoenzyme-specific π-interactions with W-dyad is colored light green. ssDNA is show as stick molecular model with filled sugars and bases. Color codes as in Figure 3B.

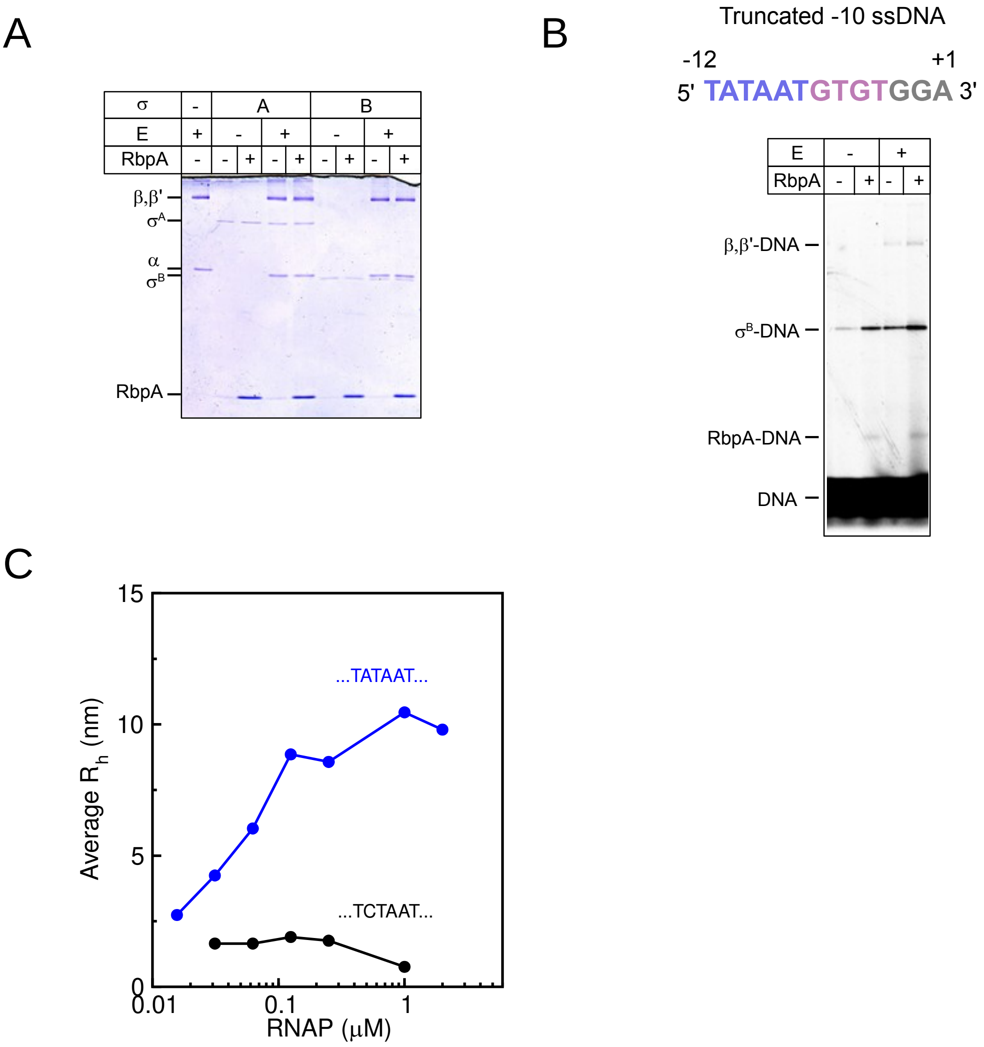


**Figure S4. Analysis of the interactions between -10 ssDNA and σ subunits**

(A) Probing of the DNA-protein interactions by formaldehyde cross-linking. Indicated combinations of MtbRNAP core, $σ^A$, $σ^B$ and RbpA were cross-linked to fluorescent -10 ssDNA and resolved on SDS-PAGE. The Coomassie blue stain is shown. (B) Probing of the DNA-protein interactions by formaldehyde cross-linking. Indicated combinations of MtbRNAP core (E), $σ^B$ and RbpA were cross-linked to a truncated version of fluorescent -10 ssDNA (shown on the top) and resolved on SDS-PAGE. (C) Measurement of -10 ssDNA (blue circles) and -11C ssDNA (black circles) binding to $Eσ^B$ by MDS. Graph shows average $R_h$ as function of protein concentration.

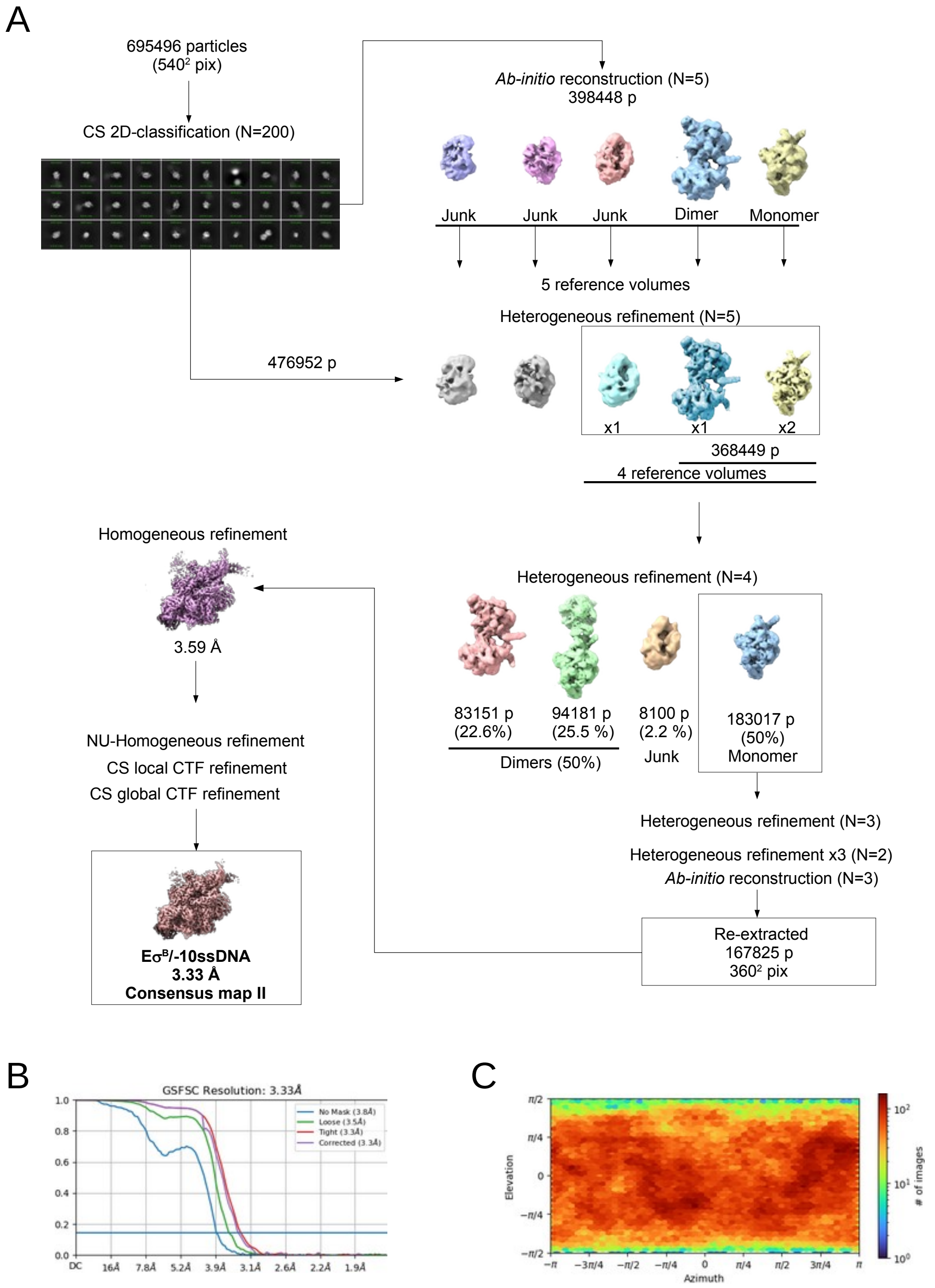


**Figure S5. Reconstruction and characterization of the consensus-II map.**
(A) cryoSPARC pipeline for consensus II map (B) Gold-standard FSC calculated for the map in cryoSPARC v3.3.2. The dotted line shows the 0.143 FSC cutoff. (C) Angular distributions for particles projections calculated in cryoSPARC and presented as a heat map.

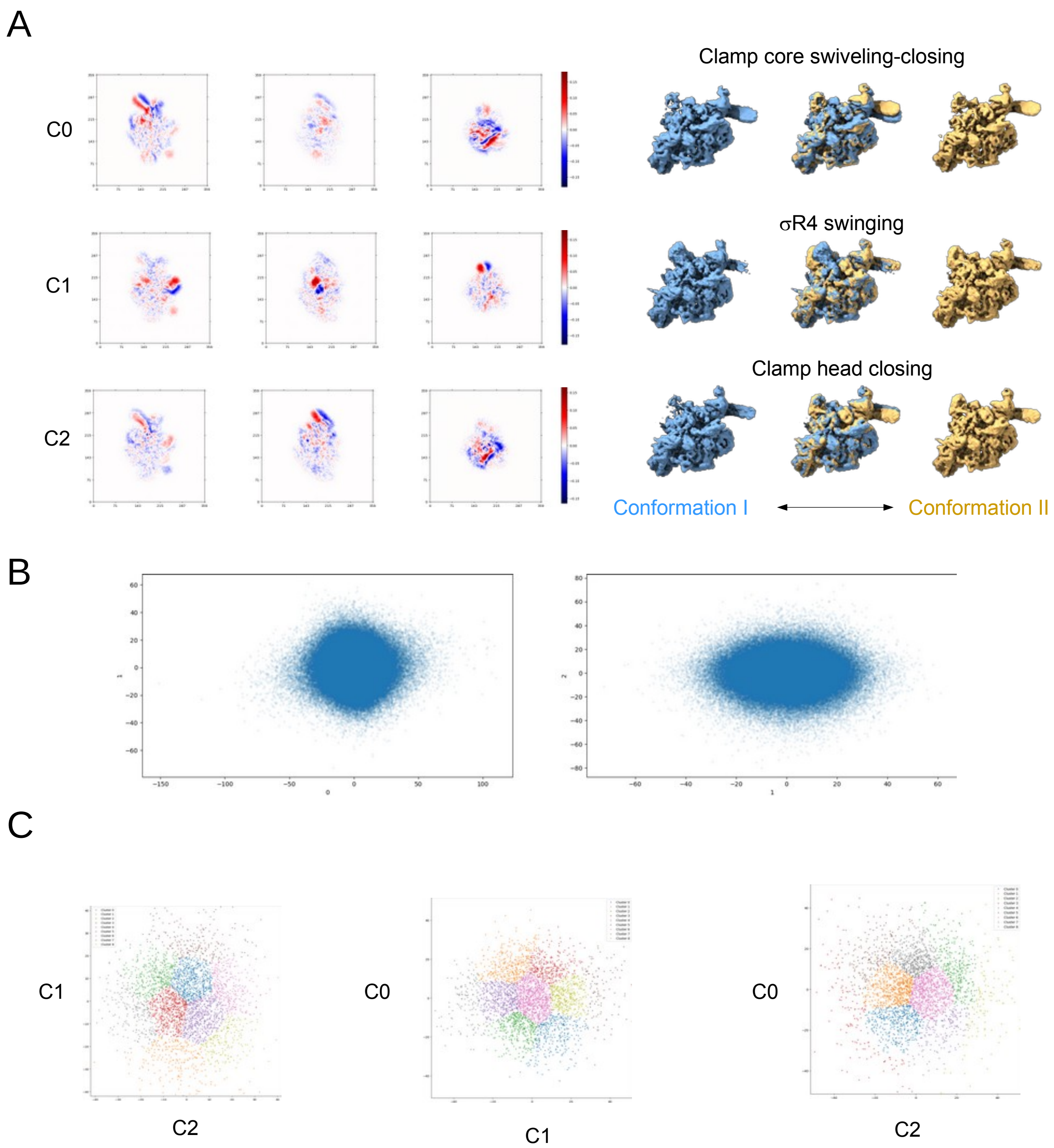


**Figure S6. Conformational heterogeneity of the E$\sigma^{B}$/-10 ssDNA complex explored by the 3D variability analysis (3DVA) in cryoSPARC**

(A) On the left: slices in x-y for three subspace directions of each reaction coordinate, C0, C1 and C3. The slices show positive (red) and negative (blue) values. On the right: 3D density maps generated in 3DVA along each variability component. Two utmost representative conformations are shown . (B) 2D-scatter plots of particles coordinates distribution over 3 components. (C) 3DVA cluster analysis of the particles latent coordinates to reveal cross-correlation between variability over each reaction coordinate.

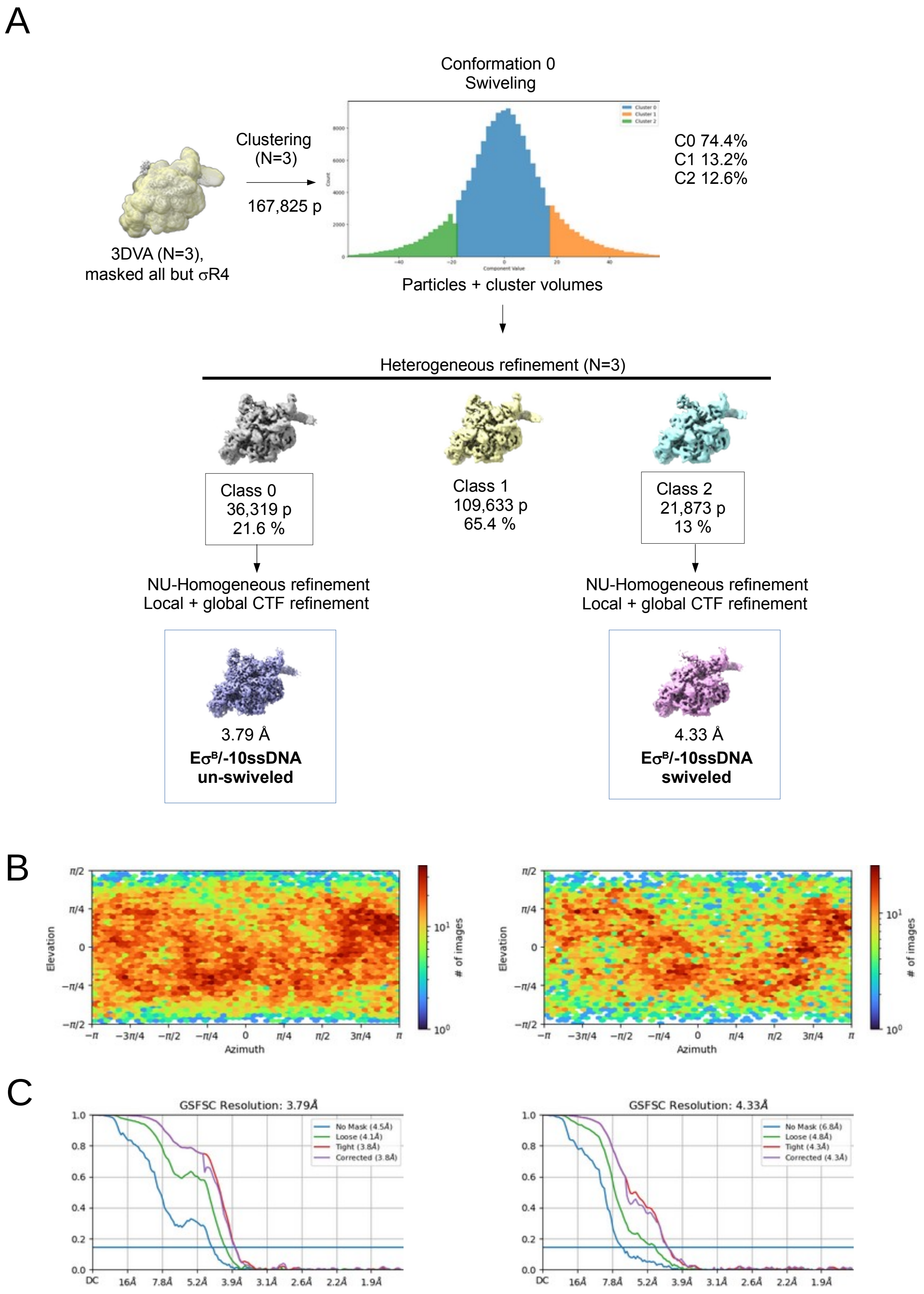


**Figure S7. Separation of the clamp conformations.**
(A) cryoSPARC pipeline for conformation 0 (left) and conformation 2 (right). 3DVA analysis was performed with particles from the consensus II map refinement job with the mask on the RNAP excluding σ subunit domain 4. (B) Angular distributions for particles projections calculated in cryoSPARC and presented as a heat map. (C) Gold-standard FSC calculated for the map in cryoSPARC v3.3.2. The dotted line shows the 0.143 FSC cutoff.

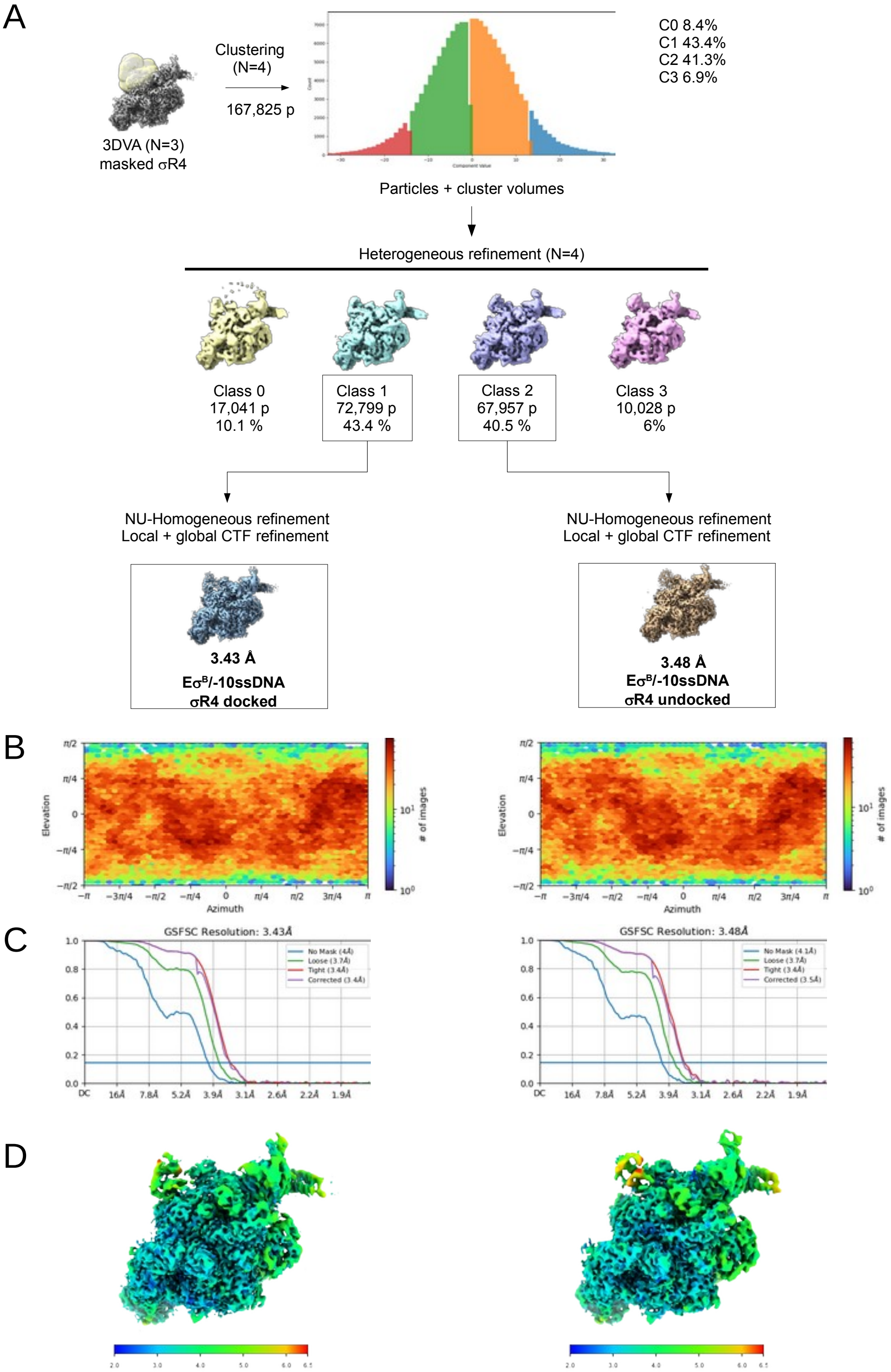


**Figure S8 Separation of the docked and undocked conformations of σR4.**
(A) cryoSPARC pipeline for docked and undocked σR4 RNAPs. 3DVA analysis was performed with particles from the consensus II map refinement job with the mask on the σ subunit region 4. (B) Angular distributions for particles projections calculated in cryoSPARC and presented as a heat map. Gold-standard FSC calculated for the map in cryoSPARC v3.3.2. The dotted line shows the 0.143 FSC cutoff. (D) Cryo-EM density maps colored according to the local resolution calculated at 0.143 FSC.
